# Regulation of antigen presentation by p38α in alveolar macrophages controls lung immunity

**DOI:** 10.64898/2026.11.26.769531

**Authors:** Monica Cubillos-Rojas, Clara Borràs-Eroles, Elisa I. Rivas, Laura Amores-Bonet, Raquel Vallejo, Elisabet Llonch, Vanessa Huaca-Manchego, Maria Papanastasatou, Stephen M. A. Forrow, Antoni Riera, Stefanie K. Wculek, Maria Tsoumakidou, Annabel F. Valledor, Angel R. Nebreda

## Abstract

The lung is constantly exposed to environmental challenges, so its immune status must be tightly regulated. Alveolar macrophages (AMs) are myeloid cells that play a key role in maintaining lung homeostasis and preventing excessive inflammatory responses. Here, we report that AMs rely on the protein kinase p38α to regulate the major histocompatibility complex class II (MHCII) expression, identifying one way in which AMs may contribute to allergic and malignant lung disease. Using genetically modified mice in which p38α is specifically deleted in myeloid cells, we show that p38α controls the antigen presentation capacity of AMs. Mechanistic studies indicate that p38α downregulates MHCII expression in AMs by restraining the expression of the transcriptional activator CIITA, which involves the p38α-regulated protein kinase MK2 and histone deacetylase 6 (HDAC6). We found that mice with p38α deleted in myeloid cells have an enhanced adaptive immune activation, which results in increased response to allergic airway inflammation but reduced tumor growth in the lung. Our results reveal that p38α signaling is required to maintain the levels of immunosuppression that preserve lung homeostasis, and show how dysregulation of this pathway may contribute to inflammatory lung diseases.

## Introduction

Lungs face the challenge of a continuous demand for gas-exchange while being exposed to foreign particles and pathogens that need to be tightly controlled to avoid diseases ^1, 2^. To balance the physiological function with the continuous exposure to potentially harmful external particles, the lungs are seeded with a variety of immune cell types that have adapted to simultaneously eliminate pathogens and restrain damage caused by inflammation that could impair gas exchange ^3^. Maintaining this delicate balance between inflammation and tolerance is critical, as excessive inflammatory responses in the lungs can cause serious life-threating diseases such as asthma, pulmonary edema, fibrosis and emphysema ^4^. In addition, lungs are the most common location for metastatic tumor growth ^5^.

Pulmonary homeostasis is preserved through the collaboration of a network of tissue-resident cells, including epithelial and endothelial cells as well as leukocytes. Lungs contain cells involved in both innate and adaptive immunity, which provide a localized tissue-specific immune protection ^6^. Among the most abundant immune cells in the lungs are alveolar macrophages (AMs), a type of specialized tissue-resident macrophage, which act as the first line of defense in the alveoli and the lung airways. Localized in the luminal side of the alveolar space, AMs perform specialized functions that keep the homeostatic and metabolic conditions required to maintain the lungs fit ^7^. Like most tissue-resident macrophages, AMs are sessile and crawl through the alveoli patrolling the zone to keep it clean from pathogens and potentially hazardous factors ^8^. Their main functions are to clear pulmonary surfactant, to phagocyte microbes and particles, and to suppress unnecessary inflammatory responses by maintaining a certain level of immunosuppression ^1, 9, 10^. Together with dendritic cells (DCs) and alveolar epithelial cells, AMs determine the threshold and quality of the immunological responses and are crucial for lung homeostasis.

AMs have a unique transcriptional signature, which is dictated by the distinctive environment conferred by the alveolar signals ^11, 12^. In homeostatic conditions, AMs are found in an immunosuppressed (M2-like) state and promote immune tolerance to prevent excessive lung inflammation. Their main function is to phagocyte foreign particles in an immunologically silent manner and to facilitate the resolution of inflammation by producing anti-inflammatory factors ^13, 14, 15^. Consistent with their immunosuppressive state, AMs are thought to have poor antigen-presentation capacity ^16^, which is supported by their low levels of major histocompatibility complex class II (MHCII) expression compared to interstitial macrophages or DCs ^17^. Antigen presentation is a fundamental process for the immune response since it connects innate and adaptive immunity. However, the contribution of AMs to antigen presentation is still unclear ^18^.

Increasing evidence supports the importance of AMs in lung inflammatory pathologies. Asthma and allergic airway inflammation are linked to the dynamics of AMs, since their strategic distribution in the lung alveoli modulates airway inflammation and influences the activation of other immune cells ^19, 20^. Although the function of AMs in tumorigenesis has been less studied than other myeloid populations such as the recruited tumor-associated macrophages or myeloid-derived suppressor cells, there is evidence implicating AMs in the development of lung tumors ^21^. It is conceivable that their distinct embryonic origin and their specialized functions in the alveolar niche provide them with specific properties to affect tumor formation. Considering their immunosuppressive and tolerogenic functions, and their presence in the lung prior to tumor formation, AMs represent perfect allies for tumoral growth.

The activation and proliferation of macrophages is regulated by signaling pathways, which upon activation by various macrophage-stimulation agents can regulate transcription factors involved in the expression of early, intermediate and late inflammatory genes ^22, 23, 24^. In particular, the mitogen-activated protein kinase (MAPK) p38α is an important regulator of immune cell functions and inflammatory responses, which controls the expression of genes involved in the innate immune response as well as in the polarization state of macrophages *in vitro* and *in vivo* ^25^.

Here, we show that p38α is crucial to maintain low expression levels of MHCII in AMs, which impacts antigen presentation and the lung adaptive immune responses. We provide evidence that the regulation of MHCII by the p38α pathway involves histone deacetylation. Experiments in mice support that p38α in AMs controls the outcome of lung inflammatory conditions such as inflammatory allergic responses and determines the growth of lung tumors. Our study sheds light on the regulation of antigen presentation in AMs and how, by controlling this process, p38α signaling impacts on lung homeostasis and disease.

## Results

### p38α downregulation affects the gene expression program of the AM population

To investigate the relevance of p38α in myeloid cells for lung immunity, we used a mouse model expressing LysM-Cre and conditional alleles of p38α, in which the exons 2 and 3 of the *Mapk14* gene encoding p38α are flanked by loxP sites, from now on referred to as p38αΔLys. The downregulation of p38α in myeloid cells was supported by immunoblotting using peritoneal macrophages (Supplementary Fig. 1a). Further analysis of myeloid cell populations sorted from the lungs of wild-type (WT) and p38αΔLys mice confirmed that the floxed exon 2 of *Mapk14* was completely deleted in AMs and partially deleted in interstitial macrophages (IMs) of p38αΔLys mouse lungs (Fig. 1a).

**Fig. 1.**
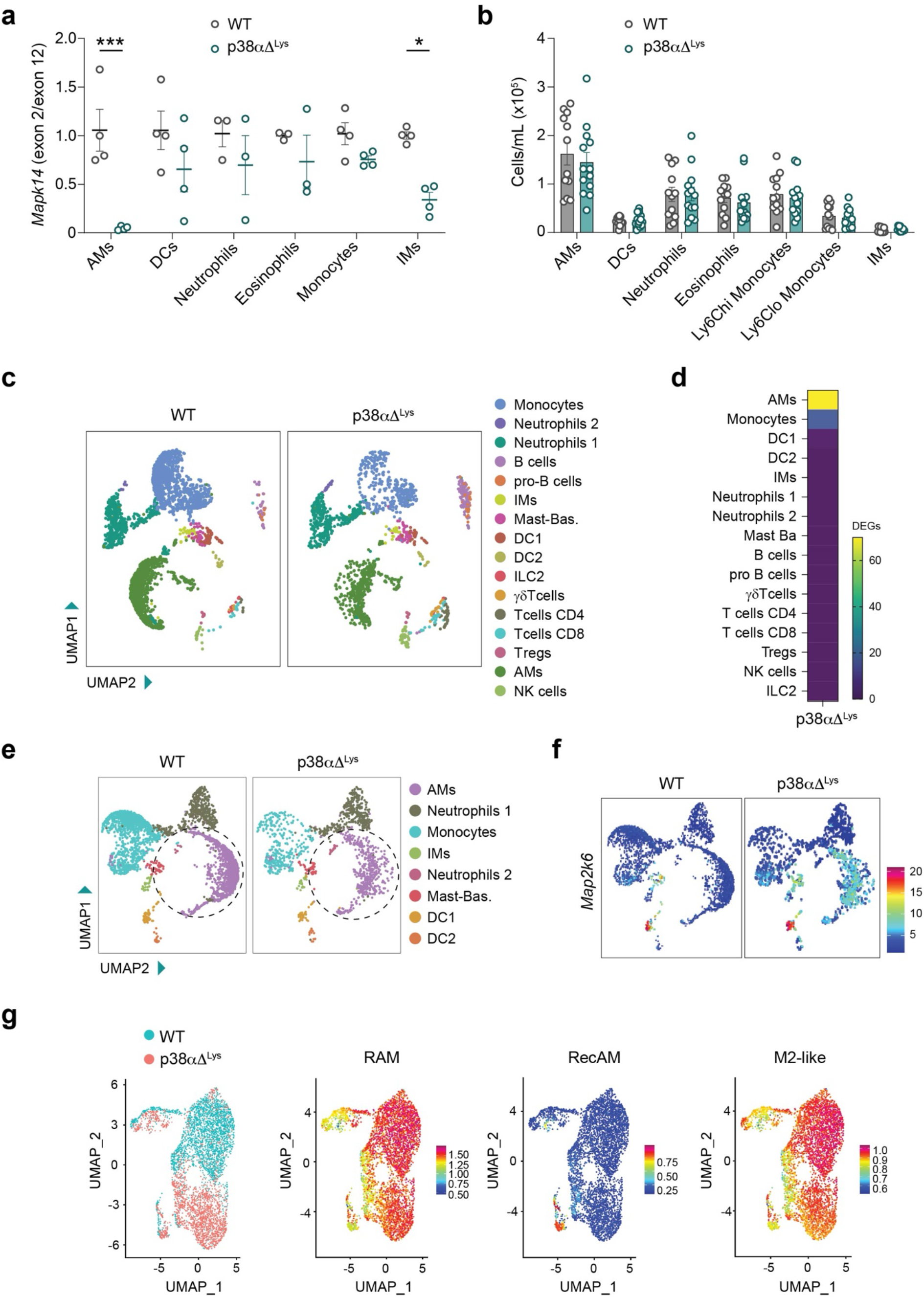
p38α downregulation in myeloid cells alters the AM population. **a.** Relative amount of the floxed exon 2 versus exon 12 (as a control) of *Mapk14* was determined by qPCR using genomic DNA of the indicated myeloid cell populations isolated from lungs of WT and p38αΔ^Lys^ mice. n=3-4. The effect of the genotype was analyzed using two-way ANOVA with a post hoc Bonferroni test. Data shown as mean±SEM. *, p<0.05; ***, p<0.0001. AMs=alveolar macrophages, DCs=dendritic cells, IMs=interstitial macrophages. **b.** Myeloid cells from lungs of WT and p38aDLys mice were analyzed by FACS. Results are indicated as the total number of cells/mL in a suspension of 350 μL coming from one lobe of the lung. WT, n=12 mice; p38aDLys, n=13 mice. **c.** UMAP representation of CD45^+^ immune cell clusters from lungs of WT and p38aDLys mice observed by scRNA-seq. Mast-Bas=mastocyte-basophil cells, ILC2=innate lymphoid cells type 2. NK=Natural killer. **d.** Heatmap representation of the differentially expressed genes (DEGs) in the indicated immune cell clusters of the scRNA-seq data. Number of DEGs in each cluster is indicated by the colored scale. **e.** UMAP representation of myeloid cell clusters in the scRNA-seq analysis of CD45^+^ cells sorted from lungs of WT and p38αΔ^Lys^ mice. **f.** Feature plot of myeloid cell clusters showing *Map2k6* expression in lungs from WT and p38aΔ^Lys^ mice. The expression levels are indicated by the colored scale. **g.** UMAP plots showing the expression levels of different reported signatures in the subcluster of AMs. The expression levels of each signature are indicated by the colored scales. Resident AMs (RAM) markers include *Mrc1* (CD206), *Itgax* (CD11c), *Siglecf* and *Siglec1*(CD169). Recruited AMs (RecAM) markers include *Cd14*, *Ly6c1* (Ly6C), *Apoe*, *Ccr5*, *Mafb* and *Sell* (L-selectin). M2-like markers include *Tgm2, Socs2, Myc, Mrc1, Klf4, Irf4, Il10, Egr2, Clec7a, Chil4, Chil3, Arg1, Alox15*.

First, we compared the numbers of myeloid and lymphoid cell populations in the lungs of WT and p38αΔLys mice using FACS analysis. We found that p38α deletion in myeloid cells did not affect the main immune cell populations in the lung, including AMs, IMs, DCs, inflammatory and circulating monocytes, eosinophils, neutrophils, CD4^+^ and CD8^+^ T cells, NK cells and B cells (Fig. 1b and Supplementary Fig. 1b). Additionally, we analyzed the baseline inflammatory status of the lungs from WT and p38αΔLys mice but observed no differences in any of the histopathological parameters evaluated (Supplementary Fig. 1c, d).

To investigate how p38α deletion in myeloid cells affects the transcriptional profile of the lung immune cells, we FACS-sorted the CD45^+^ cells of the lungs from p38αΔLys and WT mice, and performed single cell RNA-sequencing (scRNA-seq) analysis by 10X Genomics cell fractionation. This analysis identified up to 16 different immune cell populations, including AMs, IMs, type 1 and type 2 neutrophils, monocytes, type 1 and type 2 DCs, B cells, pro-B cells, γδ T cells, NK cells, CD4^+^ T cells, CD8^+^ T cells, regulatory T cells (T-regs), and group 2 innate lymphoid cells (ILC2) (Fig. 1c). In agreement with the FACS results, no differences in immune cell populations were observed between the two genotypes. However, analysis of the differentially expressed genes in each cluster showed major transcriptional changes in the AMs from p38αΔLys mice compared with WT mice, while other myeloid cell populations remained unchanged (Fig. 1d). Closer analysis of the myeloid cell sub-clustering further evidenced the importance of p38α in the gene expression programs of AMs (Fig. 1e). Feature plot representation of the expression levels of *Map2k6*, a gene that is often negatively regulated by p38α 26, revealed a higher expression of this gene in AMs compared to other myeloid cell populations from p38αΔLys mice, thereby supporting that p38α signaling is impaired in the AMs from p38αΔLys mice (Fig. 1f). Closer inspection of the AM cluster indicated a clear segregation of the gene expression profiles in WT and p38αΔLys cell populations (Fig. 1g). Alterations in the AM population cause the recruitment of circulating monocytes from the bone marrow to the alveolar space ^9^. To investigate possible differences in the origin of AMs between WT and p38αΔLys mice, we examined the expression of known markers for Resident AMs (RAMs) and Recruited AMs (RecAMs) ^27^. We found that WT and p38αΔLys AMs were both positive for RAM markers and expressed very low levels of RecAM markers (Fig. 1g). This confirms that p38α deletion does not affect the viability of the AM pool in the alveoli, and therefore does not induce the recruitment of circulating monocytes. Moreover, analysis of classical markers used to define the *in vitro* activation status of macrophages showed that both WT and p38αΔLys AMs express high levels of the M2-typical markers supporting their immunosuppressive nature (Fig. 1g). Consistent with these findings, RT-qPCR analyses confirmed high expression of RAM and M2 markers, whereas M1 associated genes were expressed at low levels in both WT and p38αΔLys AMs (Supplementary Fig. 2).

### p38α regulates crucial functions in AMs

The scRNA-seq results suggested an important role for p38α in controlling gene expression programs in AMs, which are crucial to ensure a correct lung immunity status. To further explore these effects and obtain a more mechanistic insight, we performed bulk RNA-seq analyses with AMs collected from the bronchoalveolar lavage (BAL) of WT and p38αΔLys mice. FACS analysis showed that AMs accounted for >98% of the cells in the BAL fluid of both WT and p38αΔLys mice (Fig. 2a), and both RT-qPCR and immunoblotting analyses confirmed the effective deletion of p38α in the AMs from p38αΔLys mice (Fig. 2b-c). The RNA-seq data confirmed that steady-state mRNA levels were highly different between WT and p38αΔLys AMs (Fig. 2d). We found a total of 1281 genes that were differentially expressed between WT and p38α-deficient AMs with 740 genes being upregulated and 541 genes downregulated (fold change (FC)>1.5 and false discovery rate (FDR)<0.05). This large number of changes highlights the importance of p38α in the regulation of AM functions. In fact, several genes that are involved in the homeostasis and development of AMs, such as *Car4, Bhlhe40, Bhlhe41, Krt79, Fabp4, Klf2* and *Klf4* were found differentially expressed, suggesting an important function of p38α in the biology of AMs (Supplementary Fig. 3a). We also found that some targets of PPARγ, a key signaling pathway in AMs ^28, 29, 30^, were dysregulated in the absence of p38α including *Lpl*, *Cd36*, *Cyp27a1* and *Fapb4* (Supplementary Fig. 3a, b). While these changes did not seem to affect the identity of p38α-deficient AMs (Fig. 1g), they may affect their ability to react against immune challenges.

**Fig. 2.**
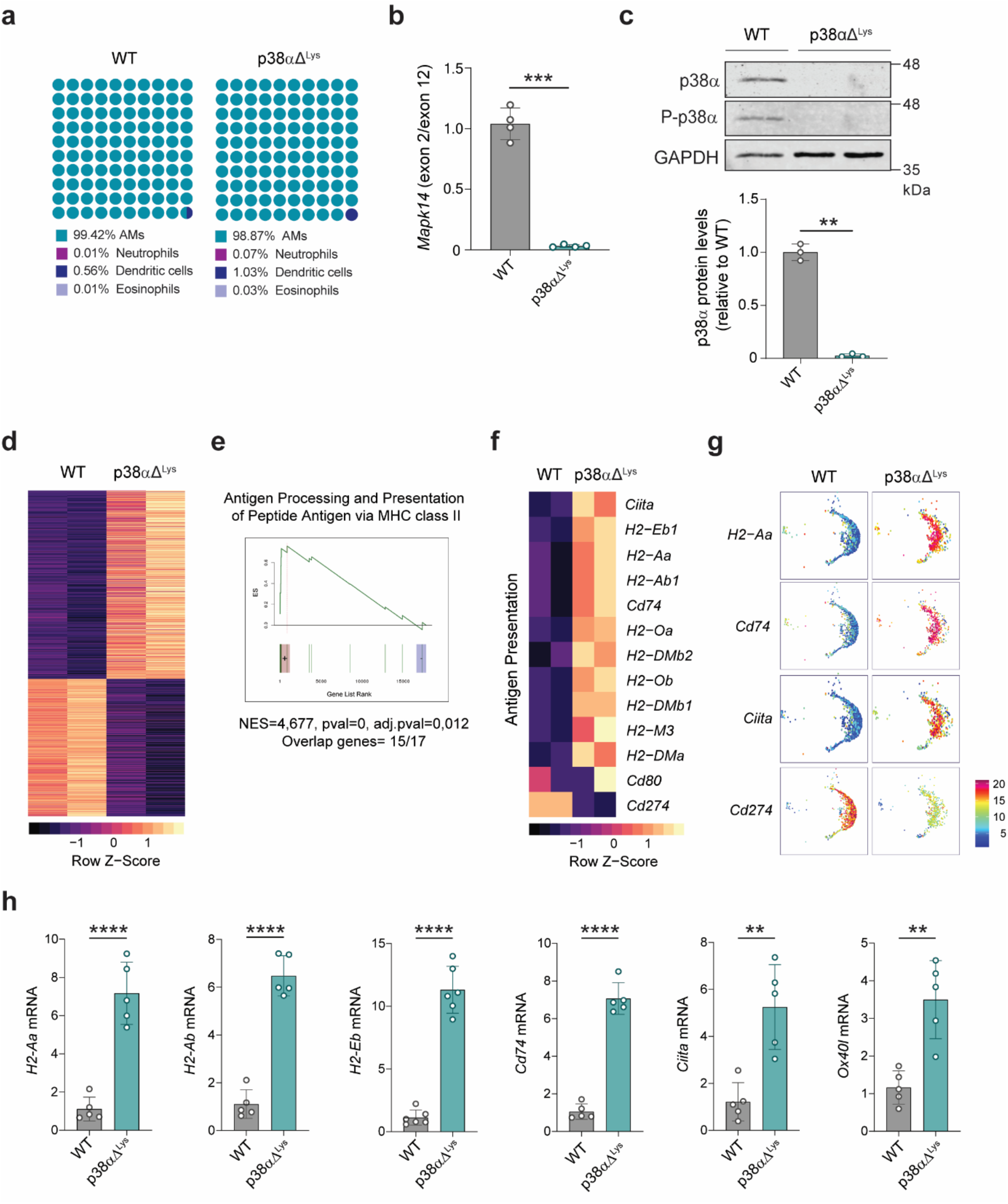
p38α regulates crucial functions in AMs. **a.** Dot plots representing the percentages of myeloid cell populations in BAL fluid from WT and p38aΔ^Lys^ mice analyzed by FACS. Each cell type is colored as shown in the legend. **b.** Relative mRNA amount of the floxed exon 2 versus exon 12 (as a control) of *Mapk14* was determined by RT-qPCR using RNA of WT and p38αΔ^Lys^ AMs extracted from BAL fluid. n=4. Data shown as mean ±SEM. Unpaired Student’s t test. ***, p<0.001. **c.** Representative images of p38α and phospho-p38α immunoblotting in AMs obtained from BAL fluid of WT and p38αΔ^Lys^ mice. The histogram represents p38α/GAPDH ratios relative to WT mice. Each dot represents a cellular lysate obtained from a different mouse (n=3). Unpaired Student’s t test. **, p<0.01. **d.** Heatmap representation of the differentially expressed genes in AMs obtained from WT and p38aDLys mice. Each line represents a pool of 2 mice. Genes were selected using fold change (FC)>1.5 and false discovery rate (FDR)<0.05 in p38aDLys relative to WT, and are presented by FC in ascending order. The relative expression levels are indicated by the colored scale. **e.** Gene Set Enrichment Analysis (GSEA) of Antigen Processing and Presentation of Peptide Antigen via MHCII (GOBP:GO0002495) comparing p38aDLys with WT AMs. ES=Enrichment Score, NES=Normalized Enrichment Score, adj.pval=adjusted pval. **f.** Heatmap representation of manually selected genes involved in antigen presentation and activation of T cells that are significantly dysregulated in the RNA-seq analysis of AMs from p38aDLys versus WT mice. The expression levels are indicated by the colored scale. p<0.05 for all the indicated genes. **g.** Feature plots of representative genes involved in antigen presentation and activation of T cells in the AMs cluster of the scRNA-seq analysis comparing p38aDLys versus WT mice. The expression levels are indicated by the colored scale. **h.** RT-qPCR analysis in AMs obtained from BAL fluid of WT and p38aDLys mice of genes involved in the antigen presentation pathway. n=5 mice from two independent experiments. Data shown as mean ±SEM. Unpaired Student’s t test. **, p<0.01; ***, p<0.001; ****, p<0.0001.

To identify potential functions regulated by p38α in AMs, we performed gene set enrichment analysis (GSEA) of the RNA-seq data using the Gene Ontology (GO) database. Interestingly, we identified antigen presentation via MHCII as one of the top upregulated pathways (Fig. 2e and Supplementary Fig. 3c). Specifically, several genes involved in antigen presentation, such as the MHCII subunits *H2-Eb1*, *H2-Aa*, *H2-Ab* or *Cd74*, were highly upregulated in p38α-deficient AMs (Fig. 2f, g). In addition, the checkpoint inhibitor *Cd274* (PD-L1) was downregulated in p38α-deficient AMs, suggesting a higher capacity to activate the adaptive immune response. RT-qPCR analyses confirmed that these genes were differentially expressed upon p38α deletion in AMs (Fig. 2h). Moreover, p38α-deficient AMs also expressed higher levels of *Ox40l*, a known activator of T effector cells (Fig. 2h). Given that MHCII and co-stimulatory molecules are associated with a more immunogenic phenotype, the predicted enhanced capacity for antigen-presentation in p38α-deficient AMs is interesting considering that these cells are expected to maintain low levels of MHCII ^17, 18^ to avoid exacerbated inflammatory responses.

The scRNA-seq analysis showed that other myeloid cell populations that have partial or no deletion of p38α expressed similar MHCII levels in both WT and p38αΔLys mice, indicating that the regulation of antigen presentation by p38α is restricted to AMs (Supplementary Fig. 3d). This was confirmed by FACS analysis of MHCII expression in IMs and DCs, the main antigen-presenting cells with highest levels of MHCII, from the lungs of WT and p38αΔLys mice (Supplementary Fig. 3e, f). Overall, our results suggest that p38α signaling limits the ability of AMs to regulate antigen presentation, which in turn likely affects their capacity to deal with lung inflammatory challenges.

### MHCII expression is regulated by p38α in AMs

We confirmed that the increased MHCII mRNA levels observed in p38αΔLys AMs correlated with a significantly higher MHCII expression at the cellular membrane (Fig. 3a). Given the exposure of AMs to specific alveolar niche factors, we investigated the regulation of MHCII by p38α in other tissue-resident macrophages. We found that peritoneal, white adipose tissue (WAT), dermal and intestinal macrophages isolated from WT and p38αΔLys mice showed similar levels of MHCII (Fig. 3b-e). Moreover, comparison of the MHCII expression profiles in different populations of tissue-resident macrophages, indicated that MHCII levels were actually downregulated in WT AMs (Fig. 3f), consistent with the idea that p38α is required to maintain low levels of MHCII expression specially in AMs but not in other specialized tissue-resident macrophages. We confirmed that the LysM-Cre recombinase was effective in all the analyzed tissue-resident macrophages, with complete p38α deletion in peritoneal macrophages and partial deletion in the remaining cell populations (Fig. 3g). Altogether, our data indicates that the negative regulation of MHCII by p38α is probably niche-specific and restricted to the alveolar space, suggesting that p38α makes a special contribution to the immunosuppressive state of the lungs. To rule out the possibility that the observed effects in AMs were caused by LysM-Cre expression in other cell types, we used an alternative Cre model under the control of the integrin alpha X (*Itgax*/*Cd11c*) promoter. We confirmed that p38α-floxed mice expressing Itgax-Cre, from now on referred to as p38αΔItgax, showed complete deletion of the p38α gene in AMs (Fig. 4a), as observed in p38αΔLys mice. We also found a significant p38α deletion in DCs, a myeloid cell population where Itgax-Cre is expected to be active. Immunoblotting confirmed the absence of p38α protein in AMs from p38αΔItgax mice (Fig. 4b). Next, we corroborated that the absence of p38α correlated with significantly increased expression of MHCII in the AMs from p38αΔItgax mice (Fig. 4c, d), but not in DCs (Fig. 4e, f). Together, these results confirm that p38α negatively regulates MHCII expression in AMs.

**Fig. 3.**
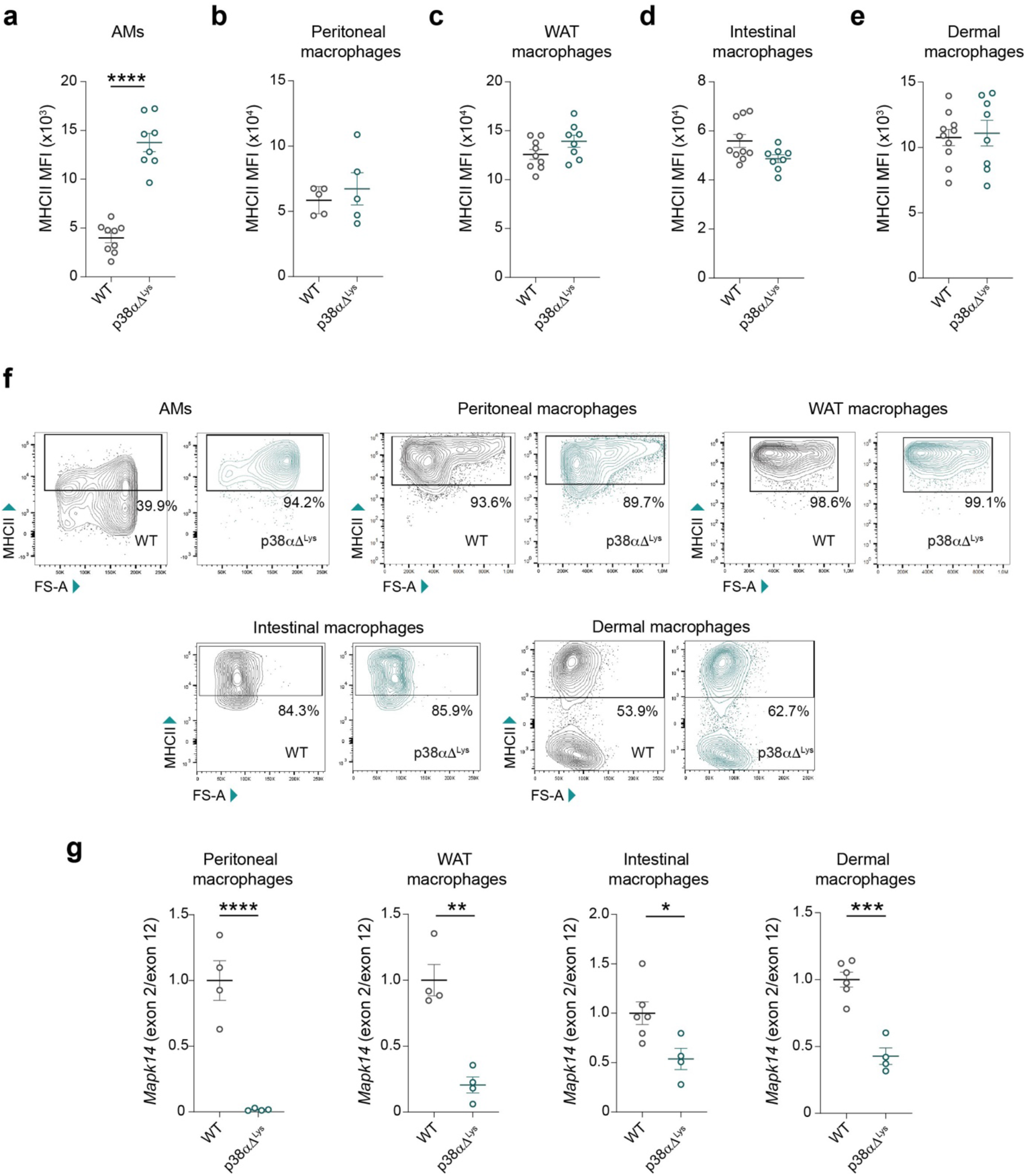
MHCII expression is regulated by p38α in AMs but not in other tissue resident macrophages. **a-e.** FACS analysis of MHCII expression in AMs (**a**), peritoneal macrophages (**b**), WAT macrophages (**c**), intestinal macrophages (**d**) and dermal macrophages (**e**) from WT and p38aDLys mice. Each dot represents cells from a different mouse. Mean Fluorescence Intensity (MFI) is shown as mean±SEM. Unpaired Student’s t test. ****, p<0.0001. **f.** Representative FACS gating plots of MHCII and Forward-Scatter (FS-A) of the indicated macrophage populations comparing WT and p38aDLys mice. The percentage of MHCII^+^ cells is indicated. **g.** Relative mRNA amount of the floxed exon 2 versus exon 12 (as a control) of *Mapk14* was determined by RT-qPCR using RNA of the indicated macrophage populations isolated from WT and p38αΔ^Lys^ mice. Each dot represents cells from a different mouse. Fold increase in p38αΔ^Lys^ macrophages was normalized relative to WT (set as 1). Data shown as mean ±SEM. Unpaired Student’s t test. *, p<0.05; **, p<0.01; ***, p<0.001; ****, p<0.0001.

**Fig. 4.**
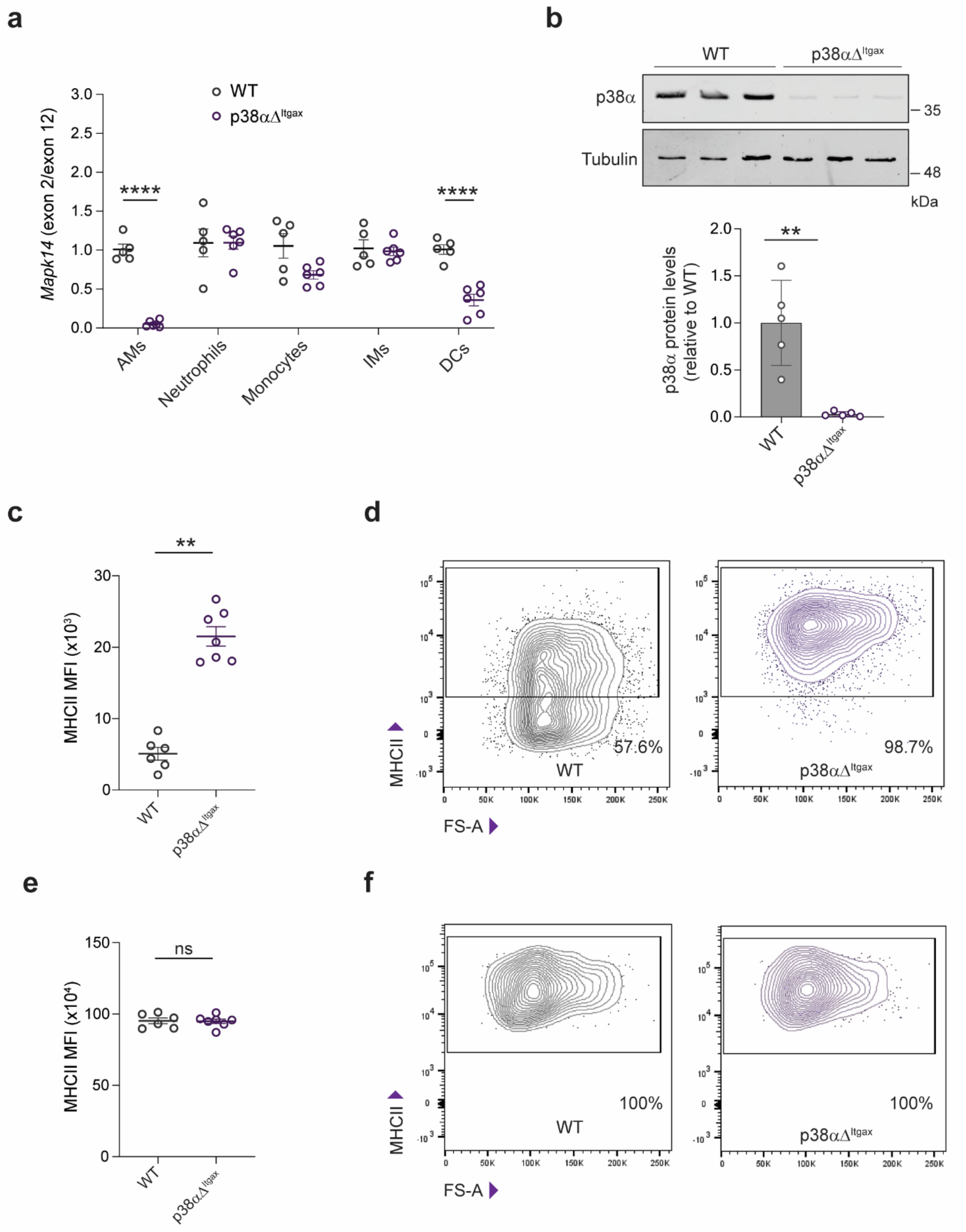
The Itgax-Cre system confirms MHCII upregulation in p38α-deleted AMs. **a.** Relative amount of the floxed exon 2 versus exon 12 (as a control) of *Mapk14* was determined by qPCR using genomic DNA of the indicated myeloid cell populations isolated from lungs of WT and p38αΔ^Itgax^. Each dot represents cells from a different mouse (n=5-6). Data shown as mean±SEM. The effect of the genotype was analyzed using two-way ANOVA with a post hoc Bonferroni test. ****, p<0.0001 **b.** Representative image of p38α immunoblotting in AMs extracted from BAL fluid of WT and p38αΔ^Itgax^ mice. The histogram represents p38α/tubulin ratios relative to WT mice. Each dot represents a cellular lysate obtained from a different mouse (n=5). Unpaired Student’s t test. **, p<0.01. **c.** Mean fluorescence intensity (MFI) quantification of MHCII expression in AMs from WT and p38αΔ^Itgax^ mice. Unpaired Student’s t test. **, p<0.01. **d.** Representative dot plots showing MHCII expression in AMs from WT and p38αΔ^Itgax^ mice. The percentage of MHCII^+^ AMs is indicated. **e.** Mean fluorescence intensity (MFI) quantification of MHCII expression in DCs from WT and p38αΔ^Itgax^ mice. **f.** Representative dot plots showing MHCII expression in DCs from WT and p38αΔ^Itgax^ mice.

To determine whether the elevated expression of MHCII in p38α-deficient AMs was accompanied by increased expression of co-stimulatory molecules required for CD4+ T cell activation, we analyzed CD86, CD80, and MHCII in AMs after stimulation with LPS or IFNγ. We found a trend toward a higher percentage of CD86^+^ cells in p38α-deficient AMs while the percentage of CD80^+^ cells remained unchanged. (Fig. 5a). Interestingly, when we analyzed the expression levels of these markers, MHCII and CD86 were both upregulated in p38α-deficient AMs following stimulation with LPS or IFNγ, and CD80 increased upon LPS stimulation (Fig. 5b). Next, to evaluate if these changes correlated with an increased capacity for CD4+ T cell activation, we performed *in vitro* T cell activation assays. Co-cultures in the presence of ecotropic murine leukemia virus (EMV) peptides ^31, 32^ showed increased proliferation of EVA CD4^+^ T cells when incubated with p38α-deficient AMs compared to WT AMs (Fig. 5c, d). Likewise, the percentage of CD4^+^ T cells expressing the activation marker CD69 increased in CD4^+^ T cells co-cultured with AMs in which p38α was downregulated (Fig. 5e, f). We next addressed whether p38α deletion in myeloid cells could change the activation profiles of the T lymphocytes *in vivo*. FACS analysis revealed a tendency for increased number of effector CD4^+^ and CD8^+^ T cells in the lungs of p38αΔLys mice compared with WT mice (Fig. 5g). We observed a similar increase in the lungs of p38αΔItgax mice (Fig. 5g). Since MHCII activates CD4^+^ T cells, the increase in effector CD8^+^ T cells is consistent with the idea that activated CD4^+^ helper T cells can contribute to the activation of CD8^+^ T cells ^33^. Taken together, these observations support that the increased MHCII levels observed in AMs upon p38α downregulation could result in higher T lymphocyte activation in the lungs of p38αΔLys mice. Given the importance of this biological process, we investigated the molecular mechanism by which p38α regulates MHCII in AMs.

**Fig. 5.**
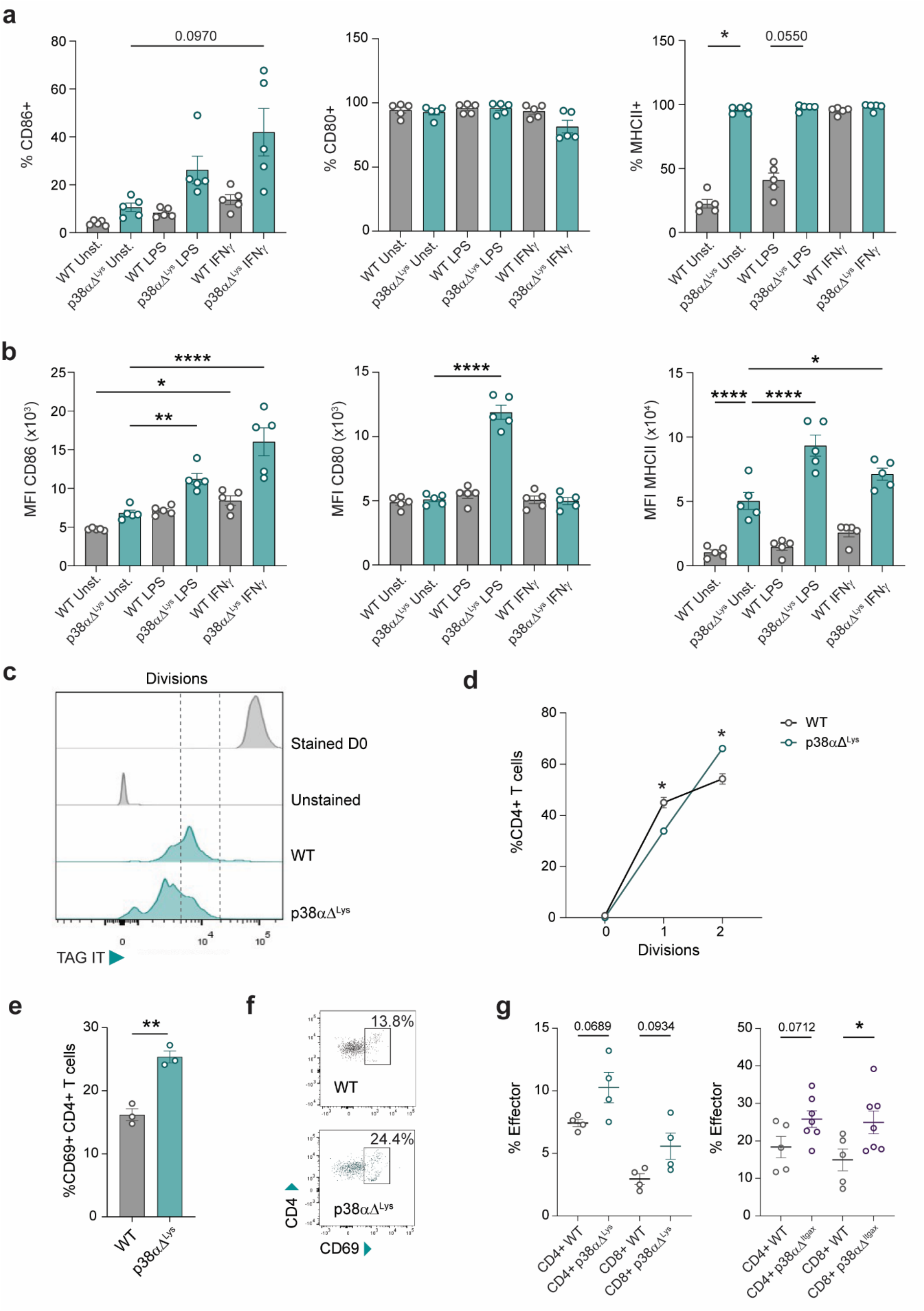
p38α regulates co-stimulatory molecule expression in AMs and T cell activation. **a.** Percentage of positive cells for CD86, CD80 and MHCII within the population of AMs obtained from BAL fluid. After isolation, cells were allowed to adhere to the plate and were stimulated with LPS (100 ng/mL) or IFNγ (50 ng/ml) for 24 h, and then were analyzed by FACS. Kruskal-Wallis with a post hoc Dunn’s test was used for multiple comparisons. * p<0.05. **b.** Mean fluorescence intensity (MFI) quantification for CD86, CD80 and MHCII in AMs from BAL fluid, stimulated as indicated in (**a**). One-way ANOVA with a post hoc Bonferroni test. Data shown as mean±SEM. *, p<0.05; **, p<0.01; ****, p<0.0001. n=5. **c.** Representative FACS plots of Tag-IT Violet fluorescent-labelled EVA CD4^+^ T cells co-cultured for 5 days with AMs from p38aDLys or WT mice previously pulsed with EMV-derived peptides. Stained day 0 (D0) and unstained controls were used to establish the proliferation gates (dashed lines). Tag-IT fluorescence dilution indicates progressive cell division. **d.** Percentages of EVA CD4^+^ T cells that have undergone 0, 1 or 2 divisions following co-cultured with either WT or p38aDLys AMs using the same culture conditions as in (**c)**. Cumulative data from three independent experiments. Each dot represents one biological replicate, corresponding to the mean of three or four technical replicates. Unpaired Student’s t test for each division. Data shown as mean±SEM. *, p<0.05. n=3. **e.** Percentages of CD69^+^ T cells within the total CD4^+^ T cell population after co-culture with WT or p38aDLys AMs. Unpaired Student’s t test. Each dot represents one biological replicate, corresponding to the mean of three or four technical replicates. Data shown as mean±SEM. **, p<0.01. n=3. **f.** Representative plots for the expression of the T cell activation marker CD69 in EVA CD4^+^ T cells co-cultured with WT or p38aDLys AMs. **g.** FACS analysis of the activation profile of T cells in lungs from WT and p38aDLys mice (left panel) or from WT and p38aDItgax mice (right panel), represented as percentage of the total CD4^+^ and CD8^+^ T cells. Effector cells were defined as CD62L^low^, CD44^hi^. Each dot represents a different mouse. Mann-Whitney test was used to compare the genotype effect in each population of T cells. Data shown as mean ±SEM from mice of two independent experiments. *, p<0.05.

### Regulation of MHCII transcription by the p38α-MK2 axis through CIITA

Unlike other tissue resident macrophages and similar to AMs, undifferentiated bone-marrow-derived macrophages (BMDMs) exhibit very low basal MHCII expression. Because BMDMs yield significantly larger cell numbers for *in vitro* experiments than AMs, we explored BMDMs as an experimental model to study the regulation of MHCII by p38α. We obtained BMDMs from WT mice, pre-incubated them with or without the p38α inhibitor PH797804, and then treated with LPS to activate the macrophages. We observed no differences in membrane expression of MHCII between BMDMs treated or not with PH797804 in the absence of LPS, and stimulation with LPS alone did not induce MHCII expression. However, macrophages pre-treated with PH797804 and subsequently stimulated with LPS showed a significant upregulation of MHCII expression (Fig. 6a). This indicates that p38α negatively regulates MHCII expression in BMDMs (Fig. 6a). Inhibition of p38α signaling by PH797804 was confirmed by the reduced phosphorylation of the downstream substrate MK2 (Fig. 6b). Consistent with these findings, LPS stimulation in the presence of the structurally distinct p38α inhibitor LY-2228820 ^34^ also increased MHCII expression. Likewise, the p38α-targeting PROTACs NR-7h and NR-11c, which induce p38α degradation ^35, 36^, produced a similar trend toward increased MHCII expression (Fig. 6c). In contrast, major histocompatibility complex class I (MHCI) levels were not affected upon p38α inhibition in BMDMs (Supplementary Fig. 4a). These results support the use of BMDMs as an experimental model to investigate the mechanisms underlying the negative regulation of MHCII expression by p38α.

**Figure 6.**
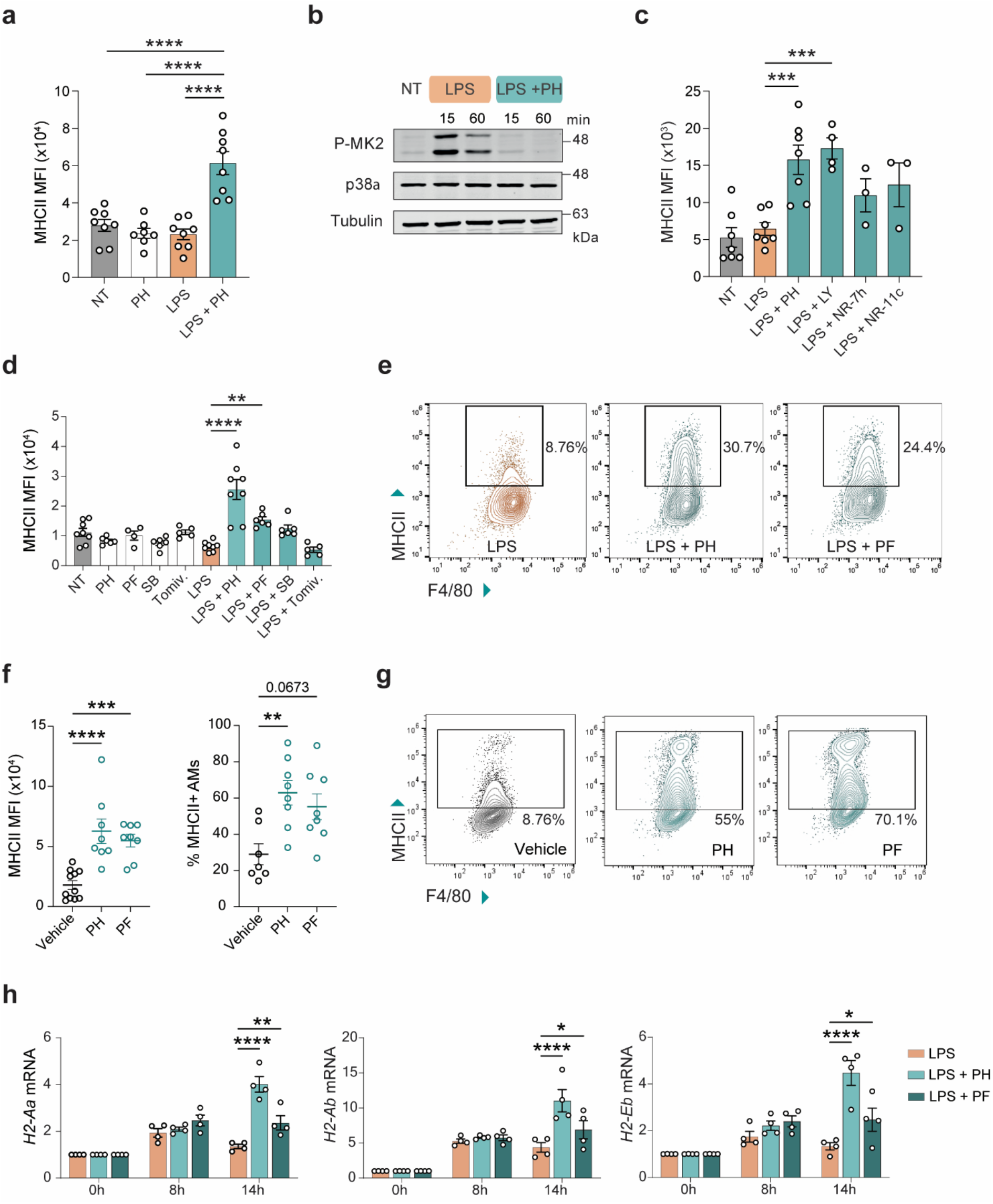
MHCII expression is regulated by the p38α-MK2 axis.a. Mean fluorescence intensity (MFI) quantification of MHCII expression in bone marrow-derived macrophages (BMDMs) from WT mice incubated with LPS for 24 h and with or without pre-treatment with the p38a inhibitor PH797804 (PH). NT=non-treated control. Each dot represents BMDMs from a different mouse. One-way ANOVA test with a post hoc Dunnett’s test. Data shown as mean ±SEM; ****, p<0.0001 referred to LPS condition. **b.** BMDMs were treated with or without PH for 2 h, stimulated with LPS for the indicated times, and then were analyzed by immunoblotting using the indicated antibodies. **c.** MFI quantification of MHCII expression in BMDMs cultured for 24 h with LPS in the presence of the p38a inhibitors PH or LY2228820 (LY) and the PROTACS NR-7h or NR-11c. Each dot represents BMDMs from a different mouse. One-way ANOVA test with a post hoc Dunnett’s test. Data shown as mean ±SEM. ***, p<0.001. p-values referred to LPS condition **d.** MFI quantification of MHCII expression in BMDMs treated for 2 h with inhibitors for p38a (PH), MK2 (PF3644022, PF), MSK1/2 (SB747651A, SB) or MNK1/2 (Tomivosertib, Tomiv) and then incubated with or without LPS for 24 h. Each dot represents BMDMs from a different mouse. One-way ANOVA test with a post hoc Dunnett’s test. Data shown as mean ±SEM. **, p<0.01; ****, p<0.0001. p-values refer to LPS condition. **e.** Representative FACS gating of MHCII expression in BMDMs treated with or without inhibitors for p38a (PH) or MK2 (PF) for 2 h and then stimulated with LPS for 24 h. **f.** Mice were intra-tracheally administered chemical inhibitors for p38a (PH), MK2 (PF), or vehicle. After three consecutive doses every 24 h, BAL was collected and MHCII levels in AMs (left panel) and the percentage of MHCII^+^ AMs (right panel) were analyzed by FACS. Each dot represents BMDMs from a different mouse. One-way ANOVA test with a post hoc Dunnett’s test was used to analyze MFI data. Kruskal-Wallis with a post hoc Dunn’s test was used for multiple comparisons of percentages data. Data represented as mean ±SEM. **, p<0.01; ***, p<0.001; ****, p<0.0001. **g.** Representative FACS plots of MHCII expression in AMs after treating the mice with three doses of the indicated p38a (PH) or MK2 (PF) inhibitors. **h.** RT-qPCR analysis of mRNAs encoding the indicated MHCII subunits in BMDMs pre-treated with or without inhibitors for p38a (PH) and MK2 (PF) for 2 h and then stimulated with LPS for the indicated times. RNA was analyzed from BMDMs obtained from 4 different mice. Two-way ANOVA test with a post hoc Dunnett’s test. Data represented as mean fold change ±SEM. *, p<0.05; **, p<0.01; ****p<0.0001. p-values refer to LPS condition.

Given that the regulation of MHCII in macrophages seems to be niche specific (Fig. 3a-e), we investigated whether typical alveolar niche factors known to modulate AM functions could signal through p38α and regulate MHCII expression. We found that GM-CSF, IL10, TGFβ and IL4 had minor effects on MHCII expression and were not affected by p38a inhibition, suggesting that the regulation of MHCII by p38α is independent on these niche factors (Supplementary Fig. 4b). In contrast, TNFa tended to increase MHCII expression, and this trend was further enhanced upon p38a inhibition (Supplementary Fig. 4b). We also tested IFNg, a typical MHCII inducer, and found that IFNg-induced MHCII upregulation was not affected by the inhibition of p38α in BMDMs (Supplementary Fig. 4c). The expression of known targets of the IFNg pathway was neither affected in AMs from p38aDLys mice (Supplementary Fig. 4d), suggesting that the regulation of MHCII by p38α is independent of IFNg signaling.

To test the possible implication of protein kinases that are directly regulated by p38α, we used chemical inhibitors. We observed that MHCII expression was not affected by the combination of MSK or MNK inhibitors with LPS (Fig. 6d). However, MK2 inhibition consistently increased the levels of MHCII in LPS-treated BMDMs to similar levels as those observed upon p38α inhibition (Fig. 6d, e). These results suggested that MK2 might contribute to the negative regulation of MHCII expression by p38α signaling. Interestingly, administration of p38α or MK2 inhibitors to mice increased both MHCII expression and the percentage of MHCII^+^ AMs *in vivo* (Fig. 6f, g), indicating that the MK2-dependent regulation of MHCII that we identified in BMDMs operates also in AMs.

Next, we analyzed the mRNA expression kinetics of different MHCII subunits and found that the *H2-Aa*, *H2-Ab* and *H2-Eb* mRNAs were all upregulated after the combined treatment of BMDMs with LPS and the p38α inhibitor in comparison with LPS alone. A smaller increase was also observed in all the MHCII subunits when LPS was combined with MK2 inhibition (Fig. 6h). This confirms that MHCII mRNA levels are upregulated in p38α-inhibited BMDMs as found in p38α-deficient AMs. The RNA-seq results (see above, Fig. 2g, h) showed the upregulation in p38α-deficient AMs of the *Ciita* gene, encoding the Class II transcriptional activator (CIITA), which is a crucial activator of MHCII gene transcription Furthermore, p38αΔLys AMs expressed higher levels than WT AMs of three different *Ciita* transcripts, namely pI, pIII and pIV (Supplementary Fig. 4e), supporting that p38α probably regulates MHCII expression through the modulation of *Ciita* transcription. In BMDMs, we observed that *Ciita* pre-mRNA levels were also upregulated upon p38α inhibition, supporting that p38α probably regulates *Ciita* transcription (Supplementary Fig. 4f). RT-qPCR analysis confirmed the upregulation of *Ciita* mRNA (Supplementary Fig. 4g), especially transcripts pIII and pIV in BMDMs treated with LPS and either p38α or MK2 inhibitors (Supplementary Fig. 4h). Similar results were observed upon TNFα stimulation (Supplementary Fig. 4i). Taken together, these results indicate that the increased expression of MHCII in p38α and MK2-inhibited macrophages is probably due to enhanced MHCII transcription mediated by the upregulation of CIITA in both AMs and BMDMs.

### MHCII regulation by p38α requires HDAC activity

Next, we analyzed if the different MHCII expression levels were maintained *ex vivo* upon culturing AMs isolated from BAL fluid (Fig. 7a). We observed that p38α-deficient AMs maintained high MHCII levels *ex vivo* for up to 8 days, suggesting that p38α regulates MHCII expression in AMs in a cell-autonomous and long-term manner, perhaps involving some type of epigenetic memory.

**Fig. 7.**
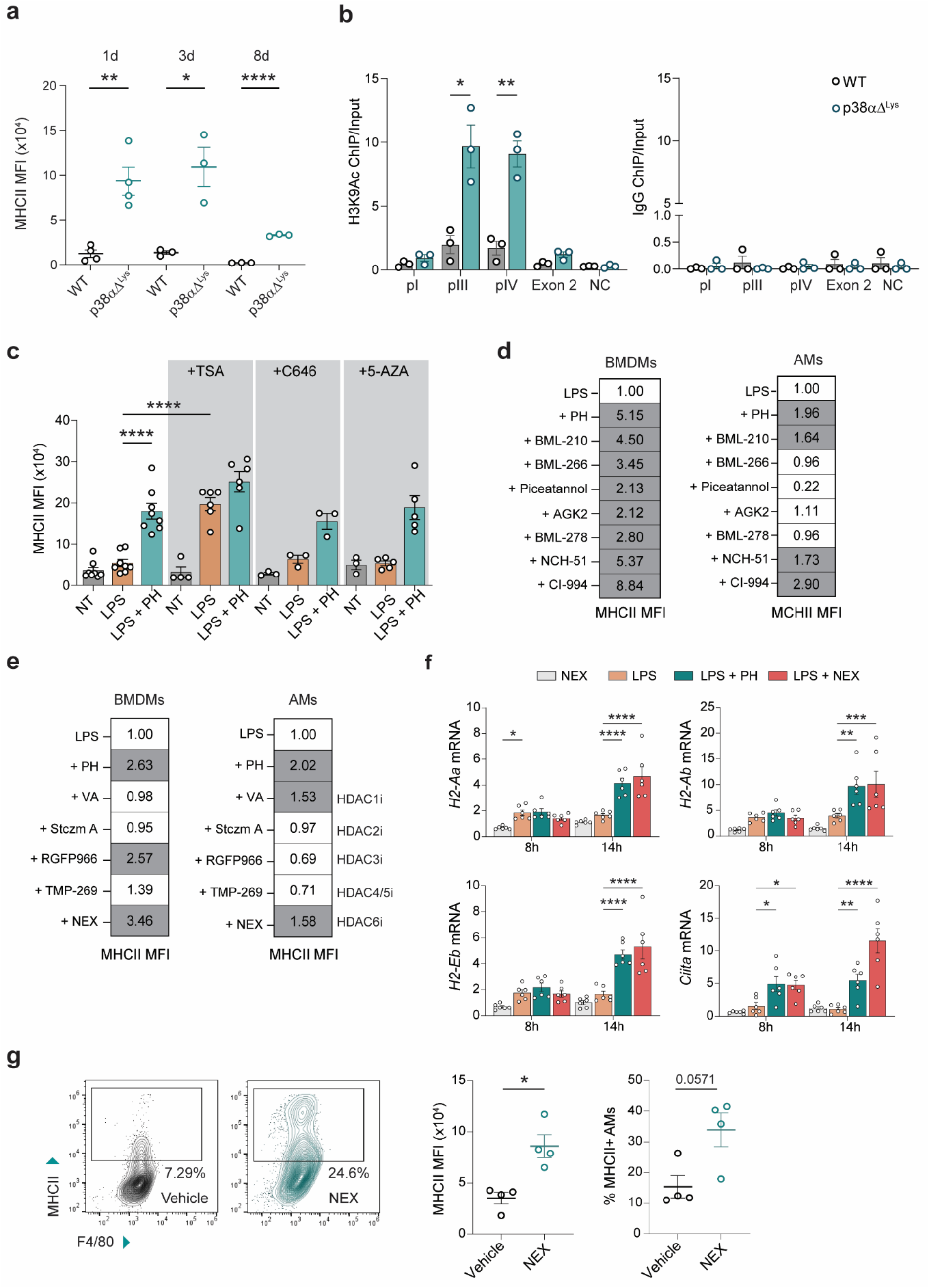
MHCII regulation by p38α involves histone deacetylase activity. **a.** FACS analysis of MHCII expression in AMs from WT and p38aDLys mice, which were cultured *ex vivo* for 1, 3 and 8 days. Each dot represents AMs from a different mouse. Data shown as mean ±SEM for each time point and genotype. Unpaired Student’s t test to compare the effect of the genotype for each time point. *, p<0.05; **, p<0.01; ****, p<0.0001. **b.** Chromatin was obtained from AMs isolated from WT and p38aΔ^Lys^ mice and was processed for Chromatin immunoprecipitation (ChIP) using antibodies for H3K9ac (left panel) or IgG control (right panel). Enrichment of H3K9ac at the indicated *Ciita* promoters, the exon2 of *Ciita* (negative control) and an intergenic region (negative genomic control=NC) was quantified by qPCR and normalized to input chromatin (% Input). Each of the three experiments were performed using a pool of AMs isolated from three mice. Two-way ANOVA test with a post hoc Bonferroni test. Data shown as mean±SEM from three independent experiments. *, p<0.05; **, p<0.01. **c.** FACS analysis of MHCII expression in BMDMs pre-treated for 2 h with the p38a inhibitor PH797804 (PH) alone or in combination with the inhibitors of histone deacetylases (TSA), the histone acetylase p300 (C646) or DNA methylation (5-AZA), and then stimulated with LPS for 24 h. NT=non-treated control. Each dot represents BMDMs from a different mouse. Two-way ANOVA test with a post hoc Dunnett’s test. Data shown as mean±SEM. ****, p<0.0001. p-values refer to LPS condition. **d.** FACS analysis of MHCII expression in BMDMs and AMs treated for 24 h with LPS in the presence of PH or the indicated epigenetic drugs (see Supplementary Table 1). All compounds were used at 20 μM, except NCH-51 and PH that were used at 2 μM. The values shown are the mean of three cellular cultures obtained from different mice. Each experiment was normalized to its LPS condition. Values >1.5 are highlighted in grey. **e.** FACS analysis of MHCII expression in BMDMs and AMs treated for 24 h with LPS in the presence of PH or the indicated HDAC inhibitors (see Supplementary Table 1). The values shown are the mean of three cellular cultures obtained from different mice. Each experiment was normalized to its LPS condition. Values >1.5 are highlighted in grey. **f.** RT-qPCR analysis of mRNAs for *Ciita* and the indicated MHCII subunits in BMDMs pre-treated with the p38a inhibitor PH and the HDAC6 inhibitor NEX for 2 h, and then stimulated with LPS for the indicated times. BMDM were obtained from four different mice. Two-way ANOVA test with a post hoc Dunnett’s test. Data shown as mean ±SEM. *, p<0.05; **, p<0.01; ***, p<0.001; ****, p<0.0001. **g.** Representative FACS plots of AMs from mice that were treated for three days with a chemical inhibitor for HDAC6 (NEX) or vehicle by intra-tracheal administration. After that, BAL fluid was collected and MHCII levels in AMs were analyzed by FACs. The left histogram represents the MFI mean ±SEM relative to vehicle-treated mice and the right histogram shows the percentage of MHCII^+^ AMs. MFI was analyzed by Unpaired Student’s t test; *, p<0.05. Mann Whitney test was used to analyze the percentage of MHCII^+^ cells.

Then we investigated if the CIITA upregulation observed in the absence of p38α was associated with an open chromatin state. Using an antibody against histone H3 lysine 9 acetylation (H3K9ac) to perform chromatin immunoprecipitation (ChIP) experiments, we observed that H3K9ac was significantly enriched at the pIII and pIV promoters of *Ciita* in p38α-deficient AMs (Fig 7b), indicating active transcription at this locus.

MHCII expression is controlled by CIITA, which is itself regulated by epigenetic modifiers ^37, 38, 39, 40^. Consequently, we explored whether epigenetic modifications, such as histone de-acetylation or DNA methylation were involved in the regulation of MHCII by p38a. To do this, we used the pan HDAC inhibitor Trichostatin A (TSA) to block class I and class II HDACs, and 5-Azacytidine (5-AZA) to inhibit DNA methylation. The histone acetyltransferase p300 inhibitor C646 was included because p300 can control *Ciita* expression ^41^ and has been described as a p38α substrate ^42^. We observed that treatment with TSA sufficed to increase MHCII levels in LPS-activated BMDMs, but no effect was observed using inhibitors of DNA methylation or p300 (Fig. 7c). These results suggest that HDACs regulate MHCII expression in BMDMs.

To extend these observations, we tested the effect of a library of compounds that target different epigenetic modifications on the expression levels of MHCII in BMDMs. We found that only compounds reported to inhibit HDACs upregulated MHCII expression in LPS-treated BMDMs to similar levels as the p38α inhibition (Supplementary Table 1).

HDAC inhibitors also consistently increased MHCII expression in LPS-stimulated AMs (Fig. 7d). The results suggest that both p38α signaling and HDACs are involved in the transcriptional regulation of MHCII in both BMDMs and AMs. Given that there are 18 different HDACs, we used more selective inhibitors to elucidate which HDAC control the expression of CIITA/MHCII by p38α, and found that the HDAC6 inhibitor Nexturastat A (NEX) significantly increased MHCII expression in both LPS-stimulated BMDM and AMs (Fig. 7e and Supplementary Fig. 5a).

HDAC6 has been linked to the regulation of gene transcription including MHCI genes in tumor cells, and in lung inflammation ^43, 44, 45, 46^. The notion that HDAC6 activity inhibits MHCII expression was further supported by the siRNA-mediated downregulation of *Hdac6*, which increased the levels of MHCII expression in LPS-treated BMDMs compared with the cells treated with control siRNA (Supplementary Fig. 5b, c). Interestingly, the *H2-Aa*, *H2-Ab* and *H2-Eb* and *Ciita* mRNAs were all upregulated upon HDAC6 inhibition (Fig. 7f) with similar kinetics as observed in cells treated with p38α or MK2 inhibitors (Fig. 6h). Importantly, we confirmed that NEX administration to mice enhanced the MHCII expression in AMs *in vivo* (Fig. 7g). These results support that HDAC6 negative regulates MHCII through *Ciita* transcription, as for p38a and MK2. HDAC6 is mainly localized in the cytoplasm, but it can be also imported into the nucleus to regulate gene expression ^47, 48^. Indeed, we found that HDAC6 localized to the nucleus in BMDMs, consistent with the idea that it can regulate gene expression in those cells (Supplementary Fig. 5d).

Taken together, our results indicate that the p38α/MK2 signaling axis downregulates MHCII expression in both BMDMs and AMs through the negative regulation of *Ciita* transcription, which probably involves HDAC6.

### p38α in AMs determines the susceptibility to lung diseases

AMs provide a supportive niche during the early stages of tumorigenesis, protecting malignant cells from the immune system ^49^. Furthermore, AMs can regulate the metastatic timing by facilitating a state of dormancy in breast cancer cells ^50^. To investigate whether the homeostatic role of p38α in AMs can affect lung tumorigenesis, we intravenously injected either the melanoma cell line B16-F10 or the lung cancer cell line LLC1 into WT and p38αΔLys mice to generate tumors in the lungs. We observed a trend toward increased effector phenotype of CD4^+^ T cells in the lungs of p38αΔLys mice with tumors (Supplementary Fig. 6a), and the MHCII expression in AMs from these animals was significantly increased (Supplementary Fig. 6b), as in the steady-state. These observations agree with the increased T cell activation observed in p38α-deficient AMs *in vitro* (see above, Fig. 5c, d). Consistently, we observed a reduced tumor burden in p38αΔLys mice compared with WT mice using both B16-F10 (Supplementary Fig. 6c) and LLC1 cells (Supplementary Fig. 6d). FACS analysis revealed no changes in the numbers of myeloid and lymphoid cell populations between the lungs of WT and p38αΔLys mice with tumors (Supplementary Fig. 6e), suggesting that the impaired tumor growth was not related to increased immune infiltration in p38αΔLys mice. However, we did not observe a reduced lung tumor burden in p38αΔItgax mice injected with B16-F10 cells compared with WT mice (Supplementary Fig. 6f). The absence of protection in p38αΔ^Itgax^ mice, despite a comparable increase of MHCII expression in AMs, could reflect the additional consequences of p38α deletion in DCs, which are efficiently targeted by Itgax-Cre. Of note, no significant differences in tumor growth were observed when B16-10 or LLC1 cells were implanted subcutaneously in WT and p38αΔLys mice (Supplementary Fig. 6g), suggesting a tissue-specific contribution of myeloid p38α signaling to tumor formation.

To complement the loss-of-function studies, we used a bacterial artificial chromosome (BAC) transgenic mouse model that has several copies of the p38α-encoding genomic region integrated in the genome, referred to as p38αBAC (Supplementary Fig. 7a). Immunoblotting analysis confirmed the upregulation of the p38α protein in AMs from p38αBAC mice (Supplementary Fig. 7b). We analyzed the MHCII levels under homeostatic conditions and observed that were similar in AMs from WT and p38αBAC mice both *in vivo* and after *ex vivo* culture (Supplementary Fig. 7c, d). Interestingly, when we analyzed mice with lungs tumors, we consistently observed decreased MHCII expression in AMs from p38αBAC mice (Supplementary Fig. 7e), consistent with the negative regulation of MHCII expression by p38α. In agreement with our previous observations, effector CD4^+^ T cells were significantly reduced in tumor-bearing lungs from p38αBAC mice compared with WT mice (Supplementary Fig. 7f). Furthermore, p38αBAC mice showed a tendency toward increased lung tumor burden compared with WT mice following tail vein injection of B16-F10 cells (Supplementary Fig. 7g).

Antigen presentation by AMs through MHCII is one of the mechanisms that contribute to the activation of the inflammatory response in the lungs, and maintenance of low levels of MHCII in AMs may help to limit excessive responses. To test the physiological relevance of MHCII downregulation by p38α in AMs, we used an inflammatory allergic model in which antigen presentation through MHCII is one of the sensitization steps that leads to eosinophilic infiltration in the airways through the Th2 response ^51^. First, we evaluated whether intranasal inoculation in WT mice with the allergen house dust mite (HDM) for 10 days, followed by analysis on the next day, induced histological changes compatible with an allergic airway response. We observed predominantly mixed-cell perivascular inflammation extended into the adjacent bronchioles (peribronchiolar inflammation). The inflammatory infiltrates were mostly located in the adventitia of affected vessels and comprised a mixed population of polymorphonuclear and mononuclear inflammatory cells. Concurrently, affected blood vessels exhibited activated endothelial cells and subendothelial polymorphonuclear cell infiltration. In addition, incipient epithelial hypertrophy and intracellular mucin accumulation were observed (Supplementary Fig. 8). When we carried out a semiquantitative assessment of inflammation across the different genotypes and treatments, we clearly observed inflammation in both genotypes upon HDM exposure in comparison to their counterpart saline controls. Interestingly, the lung inflammation was more pronounced in HDM-exposed p38αΔLys mice than in WT mice (Fig. 8a), and the same was observed when the PAS-positive bronchial epithelial area was quantified (Supplementary Fig. 9a). In addition, p38αΔLys mice exposed to HDM showed a tendency for higher IgE serum concentrations compared with HDM-treated WT mice (Supplementary Fig. 9b).

**Fig. 8.**
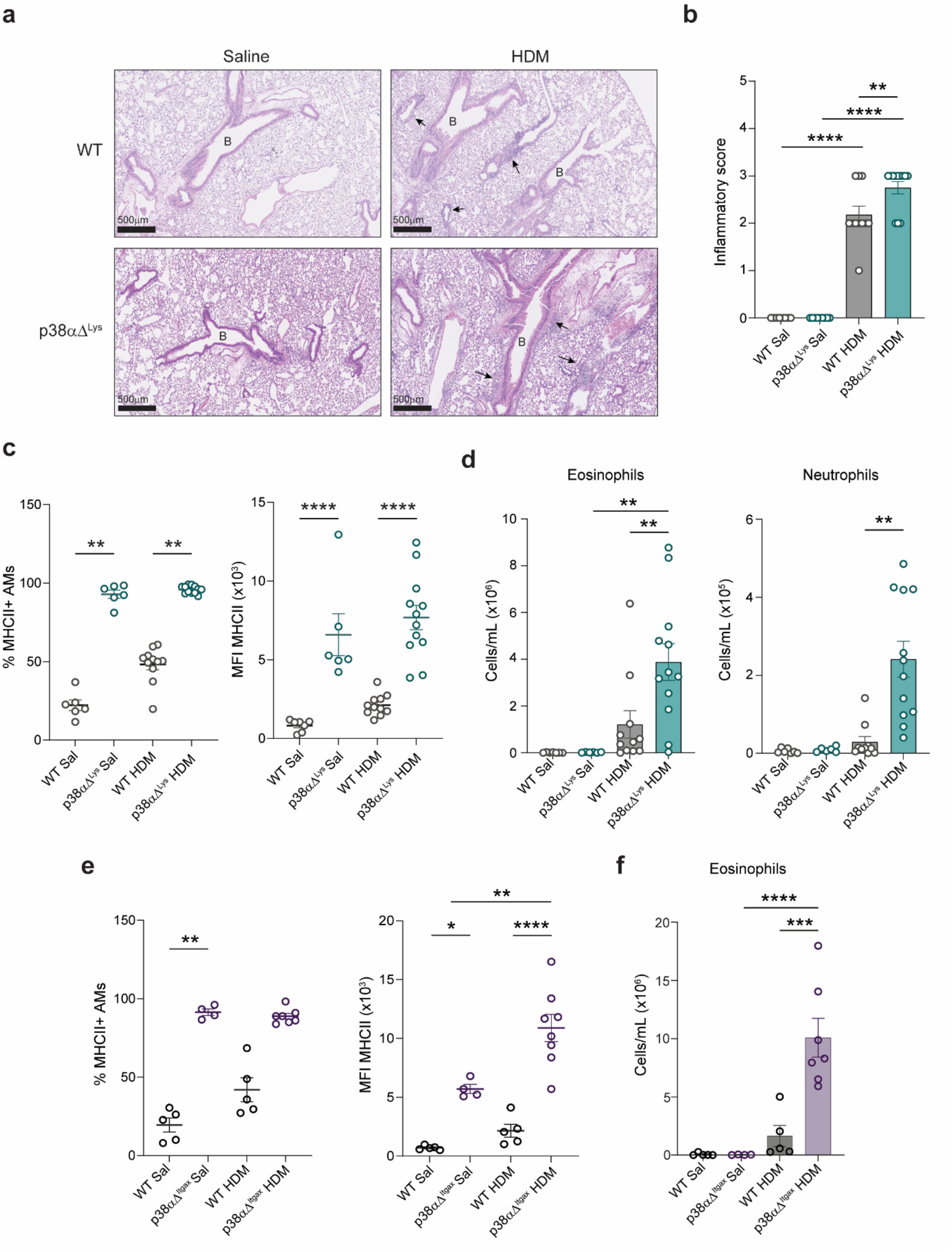
p38α in AMs determines susceptibility to inflammatory allergic responses. **a.** Representative hematoxylin and eosin (H&E) images from the lungs of WT and p38αΔLys mice exposed to house dust mite (HDM) or saline as a control. No inflammation is observed in saline-treated mice groups. WT mice show grade 2 mixed-cell inflammation, mainly perivascular, whereas p38aΔLys mice show grade 3 mixed-cell inflammation, extending around the large airways and adjacent parenchyma. Scale bars, 500 µm. Arrows indicate inflammatory infiltrates. **b.** Semiquantitative assessment of pulmonary inflammation. Severity grade was assigned in a blinded manner based on the severity and distribution of inflammation: grade 1 (minimal); grade 2 (slight); grade 3 (moderate); grade 4 (marked); grade 5 (severe). One-way ANOVA test with a post hoc Bonferroni test. Data is shown as mean±SEM; **, p<0.01. ****, p<0.0001. n=8-12 from three independent experiments. **c.** Percentage of MHCII^+^ AMs (left panel) and MHCII expression levels in AMs (right panel) from WT and p38aDLys mice after treatment with HDM or saline solution. Kruskal-Wallis with a post-hoc Dunn’s test was used to analyze percentage data. **, p<0.01. One-way ANOVA test with a post hoc Bonferroni’s test was used to analyze MFI data. Data is shown as mean ±SEM; ****, p<0.0001. n=6-12 from three independent experiments. **d.** Quantification of eosinophil and neutrophil populations in BAL fluids from WT and p38aDLys mice, which were treated with HDM or with saline solution as control. Each dot represents a different mouse. One-way ANOVA test with a post hoc Bonferroni’s test. Data is shown as mean±SEM. **, p<0.01. n=6-12 from three independent experiments. **e.** Percentage of MHCII^+^ AMs (left panel) and MHCII expression levels in AMs (right panel) from WT and p38aDItgax mice after treatment with HDM or saline solution. Kruskal-Wallis with a post-hoc Dunn’s test was used to analyze percentage data. **, p<0.01. One-way ANOVA test with a post hoc Bonferroni’s test was used for MFI data. Data is shown as mean ±SEM. *, p<0.05; **, p<0.01; ****, p<0.0001. n= 4-7 mice from two independent experiments. **f.** Quantification of eosinophils in BAL fluid from WT and p38aDItgax mice after treatment with HDM or saline solution. One-way ANOVA test with a post hoc Tukey’s test. Data is shown as mean±SEM. ***, p<0.001; ****, p<0.0001. n= 4-7 mice from two independent experiments.

We observed that exposure to HDM tended to increase both the percentage of MHCII^+^ AMs and the MHCII expression levels in AMs isolated from the BAL fluid of WT mice (Fig. 8c). Importantly, the upregulation of MHCII levels and the high percentage of MHCII^+^ AMs from p38αΔLys mice were maintained upon HDM exposure (Fig. 8c). We then examined whether this was restricted to AMs. It has been reported that HDM-exposed mice exhibit increased MHCII levels on IMs, which can contribute to allergic airway inflammation ^52^. However, in our experimental conditions, we observed no differences in the MHCII levels of IMs between WT and p38αΔLys mice exposed either to saline or to HDM (Supplementary Fig. 9c).

Furthermore, HDM sensitization increased the number of eosinophils in the BAL fluid of WT mice and this increase was substantially higher in HDM-treated p38αΔLys mice (Fig. 8d). We also observed increased numbers of neutrophils (Fig 8d), and a tendency to higher numbers of DCs and IMs in the BAL of p38αΔLys mice treated with HDM (Supplementary Fig. 8d), whereas the numbers of AMs were not affected by the allergen treatment (Supplementary Fig. 8e). Since ILC2 cells are involved in allergic responses and their activation is usually linked to Th2 responses ^53^, we checked if they were affected by HDM exposure. We observed no differences either between the saline and HDM conditions or between WT and p38αΔLys mice (Supplementary Fig. 8f). These results align with reports showing that the crosstalk between ILC2s and Th2 cells varies among mouse models, and that allergic responses can be triggered in the absence of ILC2 cell engagement ^54^.

Collectively, our data show that HDM sensitization exacerbates allergic airway inflammation in p38αΔLys mice, consistent with a role for p38α-mediated suppression of MHCII expression in AMs in limiting allergic responses.

To further support that these effects could be attributed to p38α function in AMs, we induced the allergic inflammatory response in p38αΔ^Itgax^ mice, in which p38α is deleted in CD11c-expressing cells, including AMs. We found an increased number of MHCII-positive AMs as well as higher MHCII expression in AMs upon HDM exposure, with both being more pronounced in p38αΔItgax mice than in WT mice (Fig. 8e). We also confirmed that MHCII levels did not change in IMs between treatments or genotypes (Supplementary Fig. 10a). Most of the infiltrated cells in the BAL fluid corresponded to eosinophils, which were significantly higher in HDM-exposed p38αΔItgax mice than in the treated WT mice (Fig. 8f). The concentrations of DCs and IMs were also increased in HDM-exposed p38αΔItgax mice compared with the WT mice, whereas neutrophils showed no differences between genotypes upon HDM exposure (Supplementary Fig. 10b). Histopathological evaluation revealed clear signs of inflammation following HDM exposure in both WT and p38αΔItgax mice, with a trend toward more pronounced inflammation in p38αΔ^Itgax^ mice that did not reach statistical significance (Supplementary Fig. 10c). Furthermore, serum IgE levels were significantly elevated in p38αΔItgax mice compared with WT mice upon HDM exposure (Supplementary Fig. 10d), while the frequency of ILC2 cells did not change between both groups of mice (Supplementary Fig. 10e).

Finally, we treated p38αBAC mice with HDM, and observed that the HDM-induced upregulation of MHCII in AMs from WT mice was attenuated in AMs from p38αBAC mice (Supplementary Fig. 11a). Interestingly, there was a small increase in the MHCII levels of IMs from HDM-exposed p38αBAC mice (Supplementary Fig. 11b). Analysis of the cellular composition of the BAL fluid showed that neither eosinophils nor the other myeloid cells differed significantly between HDM-exposed p38αBAC and WT mice (Supplementary Fig. 11c, d). Consistent with these results, the overall lung inflammation (Supplementary Fig. 11e) and serum IgE levels (Supplementary Fig. 11f) were similar in both genotypes.

Collectively, these data indicate that p38α acts as a negative regulator of MHCII expression in AMs. Deletion of p38α in AMs, as in p38αΔLys and p38αΔItgax mice, relieves this repression, resulting in increased basal MHCII expression. Conversely, p38α overexpression in p38αBAC mice further suppress MHCII expression to levels below those observed in WT AMs. Notably, however, this reduction in MHCII expression was not associated with an attenuated allergic response compared with WT mice, suggesting that MHCII levels in WT AMs are already sufficiently low to limit allergic inflammation.

Taken together, our findings support a role for p38a in maintaining lung immune homeostasis by restraining MHCII expression in AMs and thereby limiting excessive immune activation. While this regulatory mechanism may help maintain a low-inflammatory state under homeostatic conditions and protect against excessive inflammatory responses, such as allergic airway inflammation, it may also favor lung tumor growth by suppressing antitumor adaptive immunity. Thus, p38a signaling in AMs appears to play a dual role balancing protection against allergic inflammation with the capacity to mount effective antitumor immune responses.

## Discussion

AMs play critical roles in the regulation of lung immunity but the molecular mechanisms underlying key AM functions are not fully understood. Our results identify the protein kinase p38α in AMs as an important modulator of lung immunity. Using various approaches, we show that p38α does not control the numbers of different myeloid cell populations in the lungs, but contributes to restricting the expression of MHCII in AMs, which in turn is associated with the changes in the activation profile of T cells in the lungs. This may influence the outcome of pathological conditions, such as inflammatory allergic responses and lung tumorigenesis. Interestingly, p38α emerges as a major negative regulator of MHCII expression in AMs, but does not seem to play this function in other tissue-resident macrophages. To our knowledge, this kinase has not been previously linked to the control of MHCII in a specific type of tissue-resident macrophage. In contrast to lungs, where a certain level of immunosuppression is needed, p38α does not appear to restrain MHCII expression in peritoneal, intestinal or WAT-resident macrophages. These results not only highlight the finely-tuned regulation of MHCII in different types of antigen-presenting cells, but also the importance of the context in the study of p38α signaling.

Antigen-presenting cells provide the signals for T cell activation and it is possible that the increased number of CD4^+^ effector T cells observed in the lungs of p38αΔLys mice is related to the augmented expression of MHCII in AMs. Tissue-resident macrophages are known to locally potentiate recruited T cells after DC-priming in the lymph nodes ^18, 55^. While our results using p38αΔLys and p38αΔItgax mice, which lack p38α in myeloid and CD11c-expressing cells respectively, as well as p38αBAC mice, which overexpress it, support a connection between MHCII levels in AMs and the activation of T cells in the lungs, other immune cell populations may also act as downstream gatekeepers. For instance, T-regs ^56^ and NK cells ^57^, which are sensitive to macrophage-derived signals, might maintain the balance of T cell responses in the lung.

Our results indicate that p38α in AMs contributes to maintaining low levels of inflammation. Initial experiments connecting p38α to the production of inflammatory cytokines such as TNFα and IL6 supported the link between p38α activity and pro-inflammatory responses. However, further studies also associated p38α activity with the production of anti-inflammatory molecules like IL-10 ^58^, showing that p38α functions in the inflammatory response depend on the context and cell type ^25^. Here we show that p38α activity can limit antigen presentation in AMs, which correlates with reduced inflammatory responses in the lung. It is important to note that p38α deficient AMs exhibit a distinct activation state rather than a generalized inflammatory profile. Under steady-state conditions, their elevated MHCII expression is not accompanied by changes in the expression of inflammatory genes (*Il6*, *Il10*, *Nos2*) or baseline costimulatory molecules (CD80, CD86). Instead, p38α loss seems to prime AMs for subsequent inflammatory stimuli. Since AMs and BMDMs share key features of p38α-dependent MHCII regulation, we used BMDMs as a more tractable experimental model for mechanistic studies, bearing in mind that they do not fully recapitulate AM biology. Our ChIP assays, combined with experiments using a library of compounds that control various epigenetic modifications, support a role for the transcriptional regulation of *Ciita* and HDAC activity in controlling MHCII expression in AMs. We provide evidence that the regulation of MHCII by p38α in AMs involves the kinase MK2, which regulates *Ciita* mRNA expression probably via HDAC6. However, our results also suggest that other HDACs may regulate MHCII expression depending on the cell type, such as HDAC3 in BMDMs and HDAC1 in AMs, indicating that the two cell populations share the core pathway but not all of its components. HDACs can potentially regulate antigen presentation at different levels ^59^, and a report using the RAW 264.7 cell line provided some evidence for the implication of HDACs in CIITA regulation by p38α 60. Although these experiments were restricted to the regulation of *Ciita* pIV transcript by IFNγ, we have found that p38α also regulates the transcripts pI and pIII of C*iita*.

Establishing a causal link between p38α and HDACs is challenging, particularly because rescue experiments are difficult to design and interpret in a negatively regulated process. Nevertheless, the fact that inhibition of both p38α and HDACs resulted in very similar patterns of CIITA and MHCII upregulation supports that they could be functionally linked. We also noticed a slight additive effect on MHCII and CIITA expression upon combining HDAC inhibitors with p38α inhibition, which might be due to the HDAC inhibition affecting the transcription of MHCII independently of p38α, as histone modifications can also regulate MHCII transcription ^41^.

The proposed role of p38α in maintaining lung immunosuppression via MHCII downregulation is supported by the exacerbated inflammatory allergic response observed in p38αΔLys and p38αΔItgax mice treated with HDM. However, these results do not agree with previously published literature reporting that chemical inhibitors of p38α ameliorate inflammatory symptoms in preclinical models of asthma, chronic obstructive pulmonary disease (COPD) and acute lung injury ^61, 62, 63, 64, 65, 66, 67, 68, 69^. Because our genetically modified mice have allowed us to study the contribution of p38α in specific myeloid cells including AMs, it is likely that systemic inhibition of p38α affects other epithelial or immune cells in the lung that could differentially affect disease progression. For instance, a study modelling COPD in mice treated with LPS and cigarette smoke concluded that increased expression of p38α in type II alveolar epithelial cells was responsible for the severity of the disease ^70^. Thus, p38α inhibition can have pro- or anti-inflammatory effects depending on the cell type affected, potentially contributing to the limited efficacy of p38α inhibitors observed in clinical trials and highlighting the importance of cell-specific targeting.

Noteworthy, other studies using LysM-Cre-mediated deletion of p38α have also shown unexpected increased inflammation in models of arthritis and skin injury ^58, 71^, indicating that genetic deletion can provide mechanistic information that is not necessarily consistent with the results obtained by systemic administration of pharmacological inhibitors. Given that a major player in inflammatory allergic responses is antigen presentation by MHCII via macrophages and DCs, which is one of the early events in the Th2 inflammatory response ^72^, our findings suggest that the negative regulation of MHCII by p38α in AMs should be taken into account when testing the effect of p38α inhibitors in lung inflammatory diseases.

The consequences of MHCII upregulation are highly context dependent. Our data suggest that tight regulation of MHCII expression by p38α in AMs limits exacerbated allergic inflammatory responses. On the other hand, elevated MHCII levels on AMs are associated with protective trained immunity following viral or bacterial challenges ^73, 74^, and may restrict lung tumorigenesis by enhancing anti-tumor surveillance, as shown in our study. This context dependency highlights how the contribution of p38α to maintaining baseline immunosuppression in the lungs limits excessive immune reactions, but may also favor lung tumor formation.

Our transcriptional profiling revealed that, among the cell populations analyzed, AMs were the most prominently affected in the lungs of mice with myeloid-specific deletion of p38α. However, given the lack of AM-specific Cre lines, we cannot exclude contributions from other immune cell populations to the observed effects. For instance, p38α deficiency in cDC1s has been reported to promote Th2 responses upon HDM stimulation via the MK2-c-Fos-IL-12 axis ^75^. It should be also noted that the beneficial effects of p38α-deficient AMs could be compromised when complementary mechanisms are not fully functional, as observed in our tumor experiments using p38αΔItgax mice. Previous work has shown that p38α deficiency in DCs can impair their cross-presentation capacity and ability to prime CD8^+^ T cells ^76^, a key mechanism in the antitumor immunity. Thus, in p38αΔItgax mice, defective DC-dependent CD8^+^ T-cell priming could counterbalance the potential antitumor benefits associated with increased MHCII expression in AMs. Additionally, the absence of p38α could impair other antitumoral functions of DCs including the priming of CD4^+^T cells or NK cells ^77^.

The idea that p38α-mediated downregulation of MHCII in AMs may facilitate lung tumor formation is consistent with extensive evidence linking AMs to lung tumorigenesis ^21, 49, 78, 79, 80, 81, 82, 83, 84^. In particular, AMs have been reported to localize close to cancer cells during early lung tumor development and help tumor progression by increasing T-regs, which protect the cancer cells from CD8^+^ T cell cytotoxic activity ^49^. Accordingly, depletion of AMs was shown to reduce lung tumor development and T reg number ^49^. Our results are aligned with a recent study showing that chemical inhibition of p38α impairs bone metastasis in mice, an effect that involves myeloid cells ^85^, and with the observation that constitutively active p38α shifts AMs toward a more immunosuppressive phenotype ^86^.

Nonetheless, our study has several limitations. First, the association between increased MHCII expression and the observed allergic and tumor phenotypes remains correlative, as MHCII levels were not directly manipulated. Furthermore, given the lack of AM-specific Cre lines, we cannot fully exclude contributions from other immune cell populations. Future studies using adoptive transfer approaches or direct manipulation of MHCII expression in AMs will be important to establish their causal contribution to these phenotypes. In addition, BMDMs and AMs represent distinct macrophage populations, and BMDMs were therefore used as a complementary experimental model rather than a substitute for AMs. Although the p38α-MK2-CIITA axis functions similarly in both populations, differences in the HDACs involved suggest that the mechanisms regulating MHCII expression are not entirely conserved.

Altogether, our findings highlight the therapeutic potential of targeting AMs. Given their accessibility through aerosol-based drug delivery ^87^, and their capacity for functional reprogramming ^88, 89^, manipulating key signaling pathways in AMs may offer opportunities to reshape the lung immune microenvironment, either to limit excessive inflammation or to enhance antitumor immunity. Establishing whether p38a similarly regulates MHCII expression in human AMs will be an important step toward assessing the translational potential of this approach.

In summary, our study identifies p38α signaling as a key regulator of gene expression programs in AMs, influencing their functional properties and contributing to the maintenance of lung immune homeostasis. Importantly, by controlling MHCII expression through the transcriptional regulation of *Ciita*, p38α modulates the immunoregulatory functions of AMs, with potential implications for inflammatory lung diseases and cancer.

## Methods

### Mice

For the generation of p38αΔLys mice, floxed alleles of *Mapk14* encoding p38α 90 were combined with LysM-Cre ^91^. Animals were maintained in a C57BL/6J background and were genotyped by PCR. The efficiency of p38α deletion was checked by immunoblotting in peritoneal macrophages. The LysM-Cre transgene was always kept in heterozygosis. Littermate animals either without or with LysM-Cre but without *Mapk14*-lox alleles were used as controls. For the generation of p38αΔIgax mice, the floxed alleles of *Mapk14* were combined with Itgax-Cre mice (Tg(Itgax-cre)1-1Reiz, MGI ID:3763248) ^92^ in C57BL/6 background. Experiments were performed following the European Union, national and institutional guidelines and experimental protocols were approved by the Animal Ethics Committee of the PCB (CEEA-PCB-18-000017 and CEEA-PCB-24-000023). EVA mice were maintained in the Animal Facility of BSRC Alexander Fleming (Greece).

Mice that overexpress p38a were generated by the Mouse Mutant core facility of IRB Barcelona. The BAC clone RP24-137E7 (162,109 nt), covering the entire p38α-encoding *Mapk14* gene and approximately 74,5 kb of upstream sequence was obtained from CHORI-BACPAC (http://bacpac.chori.org). This BAC contained the *Slc26a8* gene, in the opposite orientation to the *Mapk14* coding sequence. The BAC was modified by recombineering to introduce the PiggyBac (PB) Inverted Terminal Repeat Regions (ITR’S) followed by the introduction of an Frt flanked kanamycin selection cassette to replace the unwanted sequences (approximately 57.5 kb) including the majority of the *Slc26a8* coding region but leaving approximately 20 kb of upstream *Mapk14* regulatory sequence. The Frt flanked kanamycin selection cassette was deleted by transfecting the *E.coli* containing the BAC vector with a FlpO expressing plasmid, p706-Flp (#A103, Gene Bridges). The resulting BAC, named RP24-137E7 ITR-PB Frt TP pTARBAC (102,398 nt), was purified using BACMAX DNA purification kit (#BMAX044, Epicentre Biotechnologies), dialyzed against 10 mM Tris-HCl with 0.1 mM EDTA (pH 7.4), and then mixed with the PiggyBac transposase mRNA to give a final concentration of 1 ng/μL BAC and 30 ng/μL PiggyBac mRNA. The BAC and PiggyBac mixture was injected into the pro-nucleus of 0.5 day C57B6/j mouse zygotes, which were transferred into the oviducts of 0.5 day post-coital (dpc) pseudo pregnant CD1 foster females, and allowed to develop to term. At 3 weeks of age, the offspring was genotyped for integration of the vector in the genome. Founder lines were identified by PCR genotyping at 3 weeks of age. The line was maintained by backcrossing p38aBAC/+ with WT C57BL/6 mice, and homozygous animals were generated by crossing two p38αBAC/+ mice. Quantitative PCR was used to determine the copy number in the offspring as previously reported ^93, 94^ using the primers p38α exon2 and p38α exon12 (Supplementary Table 3). Animals with two copies were considered WT, with five copies were considered heterozygous, and with more than 10 copies were deemed to be homozygous for the BAC cassette.

### Lung tumor formation

To generate lung tumors, 1.5×10^5^ B16-F10 cells or 9.5×10^5^ LLC1 cells were injected intravenously in the tail vein of 8 weeks-old female mice and allowed to grow for 21 days (B16-F10 cells) or 17-18 days (LLC1 cells). Only female mice were used in these experiments, as preliminary studies indicated that genotype-dependent differences in tumor burden were more pronounced in females. To assess pulmonary tumor burden, lungs were fixed overnight in formalin at room temperature (RT), embedded in paraffin and stained with hematoxylin and eosin (H&E). Tumor burden was quantified using the QuPath or ImageJ software as the percentage of total lung section area occupied by tumors. Three distinct lung sections were analyzed per mouse, and the mean percentage of tumor area was calculated for each animal.

### Subcutaneous tumor formation

For subcutaneous tumor formation, 1 x 10^5^ LLC1 cells or 0.5 x 10^5^ B16-F10 cells were subcutaneously injected in the right flank of 8 weeks-old female mice, and 12-14 days later, tumors were removed, weighted and measured.

### HDM-induced inflammatory allergic model

8 weeks-old mice were exposed to a purified HDM extract (Alk-Abelló) with a low LPS content (<0.5 EU/dose), as determined by the Endosafe LAL Assay (Charles River Laboratories). Mice were anesthetized with isoflurane until breathing pace was continuous, and then were intranasally inoculated with 25 µg HDM/mouse in a volume of 35 µL of 0.9% sterile physiological saline buffer, following a procedure approved by the *Generalitat de Catalunya* (DAAM8094). This procedure was repeated for 10 consecutive days as described 95, 96. Control animals were inoculated with the physiological saline buffer. After 10 days, BAL fluid was extracted and lungs were carefully collected for analyses.

### BAL

BAL was performed as previously reported ^97^. In brief, mice were euthanized using pentobarbital (200 mg/kg) and trachea was exposed. Using a 24-G shielded catheter, trachea was cannulated around 5 mm just below the larynx. The catheter was secured in place with surgical threat. A 1 mL syringe was attached to the catheter and around 7 mL of 37°C BAL buffer (2 mM EDTA, 0.5% FBS in sterile PBS) were injected and recovered in turns of 1 mL (around 800-900 μl each time). Collected BAL was kept at 37°C. Cells were pelleted at 300 xg for 5 min. If pellet was red, red blood cell (RBC) lysis buffer (5% PBS, 0.8% NH4Cl in sterile H20, filtered) was added for 1 min at RT. Cells were used for culture or for flow cytometry analyses.

### Cell line maintenance

The B16-F10 melanoma cell line and LLC1 lung cancer cell line were purchased from ATCC. The cells were cultured in high glucose DMEM medium supplemented with 10% fetal bovine serum (FBS), 1% penicillin/streptomycin (P/S) and 1% glutamine. Cells were routinely tested for *Mycoplasma* and *C.bovis*.

### BMDMs

8 weeks-old mice were sacrificed by cervical dislocation and legs were removed to isolate tibia and femur, which were kept in PBS. Bones were cleaned from skin and muscle, epiphysis of the bones was cut with sterile scissors and bones were flushed with a 5 mL of DMEM with a 25G syringe into a 60 mm Petri dish. Cells were cultured in 150 mm Petri dishes with 40 mL of warm medium (DMEM with 2% P/S + 30-40% L-929 cell (L-cell)-conditioned medium + 20% FBS) at 37°C for 6-7 days until confluent. To obtain L cell-conditioned medium, 7×10^5^ cells of the mouse fibroblast cell line L-929 (ATCC CCL 1, NCTC clone 929) were cultured in high glucose DMEM supplemented with 1% P/S, 10% FBS at 37°C and 5% CO2. Cells were grown in 150 mm plates up to confluence and 7 days later the supernatant was removed, filtered with a 0.22 μm vacuum filter and stored in aliquots at -80°C. Once thawed, the aliquots were stored at 4°C.

After differentiation for 6-7 days, BMDMs were washed with PBS and 5 mL of fresh DMEM supplemented with glutamine, P/S and 10% FBS was added to the plates (starving medium). Cells were collected with the help of a cell scrapper and were placed in non-treated 6-well plates with the starving medium for 16-18 h to allow their attachment. After this period, inhibitors were added and left for 2 h before stimulation with cytokines. Working concentrations of the inhibitors used were 2 μM PH797804 (synthesized by A. Riera’s group, IRB Barcelona), 200 nM LY2228820 (Axon Medchem 1895), 1 μM PROTACs NR-7h (Donoghue et al. 2020), and NR-11c (Cubillos-Rojas et al., 2023), 10 μM PF3644022 (Merck Life Sciences, PZ0188), 10 μM SB747651A (MedChem Express, HY-110313), 10 μM Tomivosertib (MedChem Express, HY-100022), 5 μM Nexturastat A (Selleckem, S7473), 400 μM Valproic Acid (MedChem Express, HY-10585), 10 μM Santacruzamate A (MedChem Express, HY-N0931), 10 μM RGFP966 (MedChem Express, HY-13909), 10 μM TMP269 (MedChem Express, HY-18360), 20 nM Trichostatin A (Merck Life Sciences, T1952), 10 μM 5-Aza-2’-deoxycytidine (Merck Life Sciences, A3656) and 20 μM C646 (Merck Life Sciences, 382113). Cytokines and factors were used at the following concentrations: 100 ng/ml LPS (Merck Life Sciences, L3129), 2 ng/ml TNFα (eBioscience, 14-8321-62), 20 ng/ml GM-CSF (PeproTech, 315-03), 20 ng/ml IL10 (eBioscience BMS347), 10 ng/ml IL4 (Pepro-tech, 214-14), 5 ng/ml TGFβ (PeproTech, 100-21), 50 ng/ml IFNγ (PeproTech, 315-05). PH797804 was used to inhibit p38α unless otherwise indicated. Stimulated cells were cultured for the indicated times and then were collected for analysis by FACS, RT-qPCR or immunoblotting.

For treatment with siRNA, BMDMs were electroporated using the Neon transfection system according to the manufacturer’s instructions. BMDMs (20×10^6^) were resuspended in 100 μL of buffer R (provided in the kit) and were mixed with 800 nM of control siRNA or HDAC6 siRNA (Silencer Select Pre-designed siRNAs by Thermofisher, #s67425). Cells were electroporated with 2 pulses of 1400V for 20 ms, transferred to 6 cm plates with pre-warmed medium (without antibiotics), and incubated overnight before stimulation as previously described.

### Epigenetic drug library screening

BMDMs were treated with a library of 43 different epigenetic regulators (ENZO, #BML-2836). The compounds were initially tested at 10 μM and 20 μM, but some were cytotoxic and we decreased their concentration to 2 μM, as indicated in Supplementary Table 1. After incubation for 2 h with the inhibitors, cells were stimulated with LPS (100 ng/ml) for 24 h before analysing MHCII expression by FACS.

### Isolation and culture of AMs

AMs were isolated from BAL fluid as reported ^98^. Briefly, BAL was extracted from mice (see above) and 3×10^5^-4 ×10^5^ cells were plated per well of non-treated 6-well plates containing 3 mL of pre-warmed RPMI supplemented with 1% glutamine, 1 mM pyruvate, 1% P/S and 10% FBS. After incubation for 6-18 h, medium was replaced with fresh medium containing recombinant GM-CSF at 5 ng/mL. Medium was changed every 2 days until they reached confluence approximately 7 days after seeding. To treat the cells, GM-CSF was removed the day before starting the treatments with inhibitors and LPS. Cells were collected at the indicated times.

### Isolation of peritoneal macrophages

Mice were euthanized and a small incision was performed in the abdominal skin to carefully expose the intact peritoneal area. Then, 5 mL of sterile PBS was injected in the peritoneal cavity using a 21G needle and, after a peritoneal massage, PBS was recovered. Peritoneal macrophages were isolated from peritoneal lavage by pelleting the collected PBS at 300 xg for 5 min at 4°C. Cells were seeded on non-treated 6-well plates with DMEM containing 1% P/S. After 1 h at 37°C, cells were washed twice with PBS to remove erythrocytes and non-adherent cells, and analyzed by FACS. In some cases, DMEM containing 20% FBS, 30% L-cell conditioned medium and 1% P/S was added to the cells, which were left to proliferate for 2-4 days depending on the starting cell number.

### Isolation of WAT macrophages

Mice were euthanized, the peritoneum open and the epididymal or parametrial fat pad was harvested. The tissue was cut in small pieces and transferred into a tube containing 3 ml of HBBS solution with 0.5% BSA and 4 mg/mL collagenase B (Roche, 110888150001). Tissue was incubated for 20 min at 37°C with strong shaking. Digestion was stopped by adding 10 mL of PBS with BSA and filtered using a 100 μm cell strainer (BD). The solution was centrifuged at 500 xg at 4°C, and only the small pellet fraction was kept. Pellet was resuspended in erythrocyte lysis buffer and the solution was filtered through a 40 μm cell strainer (BD). Cells were stained with antibodies and analyzed by FACS.

### Isolation of dermal macrophages

Mice were euthanized and the ears were cut close to the hairline. Ears were transferred to a 1.5 mL microcentrifuge tube containing 1 mL of digestion buffer (HBSS solution with 1% FBS, 0.04 mg/mL DNAse I, 0.25 U/ml Dispase I and 1mg/mL Collagenase A) and incubated with vigorous shaking for 2 h at 37°C. Following incubation, the cell suspension was passed through a 70 μm cell strainer (BD) and rinsed with 10 mL of FACS buffer. Samples were centrifuged at 500 xg at 4°C, the supernatant was decanted, and the cells were collected and stained for FACS analysis.

### Isolation of intestinal macrophages

Mice were euthanized, and the colons were removed and soaked in PBS to remove fat and feces. Each colon was opened longitudinally and washed in 10 mL of cold HBSS containing 2% FBS. Next, the colon was cut into small pieces and placed in a tube with 10 mL of cold HBSS with 2% FBS. Tubes were shaken vigorously and allowed to settle for 5 min before the supernatant was aspirated. The remaining tissue was incubated in a shaking water bath at 37°C for 45 min in HBSS with 2 mM EDTA. The supernatant was then removed, the tissue was washed by adding warm HBSS, and the supernatant was removed again. Following this, 10 mL of digestion buffer (RPMI-1640 supplemented with 10% FBS, 1% P/S, 1% glutamine, 0.425 mg/mL Collagenase V, 0.625 mg/mL Collagenase D, 1 mg/mL Dispase, and 30 μg/mL DNase I) was added, and the samples were incubated in a shaking water bath at 37°C for 45 min. The digestion was stopped by adding 10 mL of FACS buffer, and the cell solution was passed through a 40 μm cell strainer and centrifuged at 400 xg at 4°C. Finally, cells were collected and stained for FACS analysis.

### Antigen presentation experiments

AMs were isolated and immediately stimulated with LPS (100 ng/mL) in the presence of EMV-derived peptides for 4 h under standard culture conditions. In parallel, naïve CD4⁺ T cells were purified from the spleens of EVA mice by negative selection, excluding T-regs. Purified CD4⁺ T cells were either labeled with the proliferation-tracking dye Tag-it or left unlabeled. Following AM stimulation, CD4⁺ T cells were co-cultured with activated AMs at a 1:1 ratio in complete medium supplemented with IL-2 (10 ng/mL) and Poly:IC (1 μg/mL) for 5 days without additional stimulation. On day 5, cells were harvested and analyzed by flow cytometry. Proliferation of CD4⁺ T cells was assessed using Tag-it, and activation of unlabeled CD4⁺ T cells was evaluated based on surface CD69 expression. Data were analyzed using FlowJo (v10.10.0).

### Preparation of single cell suspensions from lungs and fluorescence-activated cell sorting (FACS) analysis

Lungs were collected and were finely minced with a scalpel. The tissue was enzymatically digested in 10 mL of protein-free DMEM containing 1 mg/mL Collagenase A (Roche, #10103586001), 0.2 mg/mL Dispase II (Sigma, #D4693) and 0.2 mg/mL DNAse I (Roche, #10104159001), during 25 min at 37°C with rotation. The enzymatic reaction was quenched by the addition of 30 mL of ice-cold 10% heat-inactivated fetal bovine serum DMEM (10% HI-FBS DMEM). Cell suspension was filtered through a 70 μm cell strainer and the remaining pieces of tissue were smashed against the filter with a 1 mL syringe plunge. The filter was washed with 10 mL of ice-cold 10% HI-FBS DMEM and the cells were pelleted at 300xg for 5 min at 4°C. Erythrocytes were lysed in Red Cell Lysis Buffer during 4 min at RT and the remaining cells were immediately washed with ice-cold 10% HI-FBS DMEM. After filtration through a 70 μm cell strainer and centrifugation, cells were resuspended in ice-cold PBS or FACS Buffer (1 mM EDTA, 4% FBS in PBS).

Prior to surface staining with the conjugated antibodies, cells were incubated 20 min at 4°C with anti-CD16/CD32 to block Fc receptor with or without Live/Dead Fixable cell dead stain kit, depending on the panel. Cells were then centrifuged and the mix of conjugated antibodies was added and incubated during 20-30 min at 4°C in the dark. The antibodies used are indicated in Supplementary Table 2. Cells were washed once with FACS buffer and resuspended in FACS buffer with or without DAPI depending on the panel.

FACS analysis and cell separation were performed in a Beckman Gallios Flow Cytometer or in a Beckton Dickinson FACS Aria Fusion flow cytometer. Data was analyzed using the FlowJo software. Immune cell populations were defined as described in Supplementary Fig. 12 and 13.

### Treatment of mice with chemical inhibitors

For intratracheal administration, mice were anesthetized with isoflurane and secured in a supine position on an inclined intubation platform. The tongue was gently extended to visualize the vocal cords, and a catheter was inserted through the trachea. A volume of 50 μL of each chemical inhibitor was delivered into the lungs daily for 3 consecutive days. The inhibitor stock solution was prepared at 3.75 mg/mL in PBS containing 5% DMSO, 0.5% methylcellulose and 0.025% Tween-20, and sonicated for 20 min. Control mice received the vehicle solution alone. The day after the third dose, BAL was collected and analyzed by FACS.

### Quantification of IgE in serum

Blood samples were collected via intracardiac puncture. Serum was recovered after allowing the blood to clot, followed by centrifugation at 1000 xg for 20 min at 4°C. Total IgE levels were analyzed by ELISA using a commercial kit (EM20006, FineTest) according to the manufactures’s instructions. Absorbance was measured at 450 nm the IgE concentration were calculated by interpolation from a standard curve.

### Lung histological analysis and assessment of pulmonary infiltration

Lungs were fixed in 10% neutral buffered formalin (NBF) at RT, embedded in paraffin and cut into 10 μm-thick sections. Three sections per lung were then stained for H&E and analyzed.

For the semiquantitative assessment of inflammation, all inflammatory changes associated with the HDM model were integrated into a single overall parameter: inflammation, vascular/perivascular, mixed cells. Lesion severity was graded on a five-point scale from Grade 1 to Grade 5 (minimal, slight, moderate, marked, and severe). The severity grade was assigned based on the overall severity and distribution of inflammation, considering the number and caliber of affected blood vessels, the extent of inflammatory infiltration into adjacent bronchiolar and peribronchiolar areas. Diagnostic terminology and lesion classification were based on international guidelines from the International Harmonization of Nomenclature and Diagnostic Criteria (INHAND) and the Society of Toxicologic Pathology (STP).

To analyze mucus production, lung sections were stained with periodic acid-Schiff (PAS). For each selected bronchus, the total airway wall was manually delineated as the Region of Interest (ROI). The PAS-positive area within the ROI was quantified, and the data were expressed as the percentage of PAS-positive area relative to the total bronchial area.

### Analysis of p38a deletion

Genomic DNA was isolated by heating the cell pellet at 95°C in 25 mM NaOH and 0.2 mM EDTA for 10 min. The reaction was stopped by adding an equal volume of Tris-HCl 40 mM, and analyzed by qPCR to determine the levels of exon2 relative to exon12 of *Mapk14* using the primers listed in Supplementary Table 3 and the Sybr Select Master Mix for CFX (Life Technologies).

### RNA-seq of AMs

AMs were extracted from BAL, pelleted at 350 xg for 5 min at 4°C and resuspended in 350 μL of Trizol. RNA isolation was performed with the PureLink RNA minikit. Total RNA extractions were quantified with the Qubit RNA Hs Assay kit (Invitrogen), and RNA integrity assessed with the Bioanalyzer 2100 RNA Pico assay (Agilent). Four mice per condition were pooled into 2 biological replicates of each condition.

Libraries for RNA-seq were prepared at the Functional Genomics core facility of IRB Barcelona. Briefly, mRNA was isolated from 140 ng of total RNA using the kit NEBNext Poly(A) mRNA Magnetic Isolation Module (New England Biolabs). RNA-seq libraries were prepared from the purified mRNA using the NEBNext Ultra II RNA Library Prep Kit for Illumina (New England Biolabs). Twelve cycles of PCR amplification were applied to all libraries. The final libraries were quantified using the Qubit dsDNA HS assay (Invitrogen) and quality controlled with the Bioanalyzer 2100 DNA HS assay (Agilent). An equimolar pool was prepared with the nine libraries and submitted for sequencing at the Centre for Genomic Regulation (Barcelona). A final qPCR quality control was performed before sequencing in one lane of an Illumina HiSeq2500. Sequencing output was 294 Million 50-bp single-end reads and a minimum of 29 million reads were obtained for all libraries.

Bioinformatics analyses were performed by the Biostatistics/Bioinformatics core facility of IRB Barcelona. In brief, RNA-seq reads were aligned to the mouse genome version mm10 using STAR (v.2.5.2b) (Dobin et al., 2013). SAM files were converted to BAM files and sorted using sambamba (v0.7.1) (Tarasov et al., 2015). The count matrix was generated with Rsubread with the built-in annotation for mm10 (Liao et al., 2019). DESEq2 (v1.30.1) was used for differential expression analysis with fold change shrinkage as implemented in the lfcShrink function (Love et al., 2014). Functional enrichment analysis was performed over gene sets defined in the Molecular Signatures Database (MSigDB) hallmark gene set collection, Gene Ontology database, and KEGG. The rotation-based approach for enrichment implemented in the R package limma was used to represent the null distribution (D. Wu et al., 2010). The max-mean enrichment statistic, under restandardization, was considered for competitive testing (Efron & Tibshirani, 2007).

### Single cell-RNA-seq of CD45^+^ cells

Lungs were collected as described above and CD45^+^ cells were sorted using an Aria Sorter. We used a pool of four mice for each genotype.

For single-cell transcriptome analysis, cell suspensions were processed at the Functional Genomics core facility of IRB Barcelona. Briefly, the cell concentration of each sample was adjusted to approximately 1,000 cells/μl. Cell partition into GEMs was performed using Chip G and the 3’ v3.1 chemistry (10X Genomics). Barcoded cDNA was amplified for eleven cycles, quantified, and quality controlled on the Bioanalyzer 2100 using a high sensitivity DNA assay (Agilent). Libraries were generated using only 10 μl (25%) of the total cDNA (40 μl), strictly following the manufacturer instructions. 13 amplification cycles were performed for each library. Purified libraries were quality controlled on the Bioanalyzer 2100 and quantified. A 20 nM equimolar pool was prepared and a final qPCR validated libraries before cluster generation. Illumina sequencing was performed at Centro Nacional de Análisis Genómico (Barcelona) using a NovaSeq6000 S2 with an asymmetric strategy (28+90) to reach 150 Gbp. Sequencing output was 1,589 million paired-end reads and a minimum of 233 million reads were obtained for all libraries.

Bioinformatics analyses were performed by the Biostatistics/Bioinformatics Facility of IRB Barcelona. Briefly, Chromium single cell RNA sequencing reads were aligned to the reference transcriptome (refdata-gex-mm10-2020-A) with CellRanger (v4.0.0) (Zheng et al., 2017). The count utility was used with default options to quantify gene expression. The subsequent processing steps and analysis were performed with Seurat package (v4.0.3) ^99, 100, 101, 102^. Cells having <20% mitochondrial read content and >500 UMIs were considered for downstream analyses. Ribosomal reads were removed. The proportion of mitochondrial reads was regressed out during the normalization and variance stabilization of raw counts, which was performed with the sctransform method ^103^. SCT transformed counts were further imputed and smoothed with MAGIC (v.2.0.3) (van Dijk et al., 2018). Cell types were annotated with singleR using as reference the ImmGen expression dataset from celldex 104. The mitochondrial and UMI content thresholds were manually refined by inspecting their distribution in each cell type. A threshold of 1000 UMIs was set for Neutrophils, T cells, NK cells, B cells, ILC, NKT, Endothelial cells, Tgd, proB cells, Basophils, Stromal cells and Eosinophils. A threshold of 1500, 2000, 3000, 4000, and 4500 was set for Monocytes, Mast cells, Macrophages, Fibroblasts, and DCs, respectively. A threshold of 5000 was set for Stem cells and Epithelial cells. The threshold of mitochondrial content ranged from 10 to 20% for all cell types. Cells that passed this filtering step were renormalized, imputed/smoothed with MAGIC and classified again with singleR. The first 10 principal components were used to obtain the Uniform Manifold Approximation and Projection (UMAP) for visualization purposes. Cells were assigned to clusters using FindClusters Seurat function (resolution = 1.2). A focused analysis was performed on the lymphoid and myeloid lineages separately. The expression of marker gene signatures ^105^ was summarized by taking the average MAGIC score of its constituent genes and then used to refine the cell type annotation manually. Pseudobulk differential expression analysis was performed for each cell type using DESEq2 with lfcShrink fold change shrinkage, as described above ^106^. Functional enrichment analysis was performed over gene sets defined in the Molecular Signatures Database (MSigDB) hallmark gene set collection, Gene Ontology database, and KEGG. The rotation-based approach for enrichment implemented in the R package limma was used to represent the null distribution ^107^. The max-mean enrichment statistic, under restandardization, was considered for competitive testing ^108^.

### RT-qPCR analysis

For RNA extraction, cells were washed with PBS and collected in 300-500 μl Trizol. 100 μL of chloroform were added and tubes were centrifuged at 15000 xg at RT for 10 min. The fraction with less density was transferred into new tubes. After adding 200 μl of 70% ethanol, the RNA extraction was followed using the PureLink RNA mini kit. DNAse treatment was performed using on-column DNase treatment following manufacturer’s instructions. RNA purity and concentration were determined by measuring absorbance at 260 nm and 280 nm using a NanoDrop 2000 spectrophotometer (Thermo Scientific). For sorted cells, RNA was extracted using the PureLink RNA micro kit following the manufacturer’s instructions.

cDNA was obtained from 150 ng to 1 μg of purified RNA. First, RNA was incubated with random primers and dNTP at 65°C for 5 min and ramped down to 4°C in a BioRAD apparatus (PTC0220). Samples were put on ice and a mixture of RNAsin and SuperScript IV reverse transcriptase was added. The PCR was run with the following program: 23°C for 10 min, 55°C for 10 min and 80°C for 10 min. Samples were stored for a maximum of 1 week at 4°C or for longer at -20°C.

For RT-qPCR, 3-15 ng of cDNA were mixed with 5 μl of SYBR green, 0.25 μL of each primer and up to 10 μL of ddH2O. Primers are described in Supplementary Table 3. For pre-mRNA analysis, primers were designed within the pre-exon 2 intronic region using the USCS database (Zeisel et al., 2013).The mix was incubated in a Quant6 Flex PCR apparatus (Thermofisher) with the following program: an initial step of 50°C for 2 min and 95°C for 10 min; 40 cycles of 95°C for 15 sec and 60°C for 1 min; and a final step of 95°C for 15 sec, 60°C for 1 min and 95°C for 15 sec.

Samples were analyzed in triplicates and normalized to HPRT and/or GAPDH. Analysis was done using the ΔCt method.

### Chromatin immunoprecipitation

AMs from three mice were pooled for each reaction and chromatin immunoprecipitation (ChIP) was performed using a commercial ChIP assay kit (17-295, Millipore) according to the manufacturer’s instructions. Immunoprecipitation reactions were carried out using either 5 μg of anti-H3K9ac antibody (39017, Active Motif) or normal rabbit IgG (2729, Cell Signaling) as a negative control. For DNA recovery, one volume of phenol:chloroform:isoamyl alcohol (25:24:1) was added to the enzymatic digests, and samples were vortexed for 20 sec. Phases were separated by centrifugation at 16000 xg for 5 min at RT. The upper aqueous phase was transferred to a clean tube, and DNA was precipitated overnight at −20°C by adding 20 μg glycogen, 0.5 volumes of 7.5M NH4OAc and 2.5 volumes of 100% ethanol. The precipitated DNA was pelleted by centrifugation at 16000 xg for 30 min at 4°C. The resulting pellet was washed with 70% ethanol, air dried at RT, resuspended in nuclease-free water, and quantified using a Nanodrop. For qPCR analysis, reactions were performed in triplicate using 12 ng of DNA per well. Primer sequences are described in Supplementary Table 3.

### Immunoblotting

Cells were washed twice with PBS, placed on ice and lysed with RIPA buffer (1% NP40, 0.5% sodium deoxycholate, 0.1% SDS, 50 mM Tris-HCl, pH 7.5, 150 mM NaCl, 5 mM EDTA, 5 mM EGTA, 20 mM sodium fluoride, 1 mM PMSF, 1 mM sodium orthovanadate, 2.5 mM benzamidine, 10 μg/mL pepstatin A, 1 μM mycrocystin, 10 μg/mL leupeptin and 10 μg/mL aprotinin) using a scraper while maintaining the plates on ice. After incubation on ice for 10 min, the cell lysates were spun at maximum speed for 15 min at 4°C, and the supernatant was collected and either kept at -80°C or used for protein quantification. Protein concentration was estimated using the RC DC protein assay kit I, and concentrations were calculated using a BSA standard curve, which was prepared and quantified in every experiment.

Proteins (15-40 μg) were mixed with loading buffer, boiled for 5 min at 95°C, and separated by NuPAGE gradient gels 3-8% (Invitrogen) or SDS-PAGE gels. After electrophoresis, proteins were transferred onto a nitrocellulose membrane using a wet transfer system (Bio-Rad). The membrane was blocked for 1 h with 5% non-fat milk in PBS+0.01% Tween at RT. Primary antibodies were diluted with 5% BSA in PBS-0.01% Tween and incubated overnight at 4°C. Membranes were washed three times in PBS+0.01% Tween and incubated for 1 h at RT with the secondary antibody diluted in the same buffer containing 5% non-fat milk. Finally, membranes were washed three times with PBS+0.01% Tween. The antibodies used for immunoblotting were anti-p38α (1:500; Cell Signaling, 9218S), anti-HDAC6 (1:500; Cell Signaling, 7612S), anti-phospho-T180/Y182 p38α (1:1000; Cell Signaling, 4631S), anti-phospho-T334 MK2 (1:500; Cell Signaling, 3007S), anti-Lamin A/C (1:500; Santa Cruz Biotechnology, sc-20681), anti-Tubulin (1:10.000; Sigma, T9026), anti-GAPDH (1:10.000; Sigma, G8795). Proteins were detected using the Odyssey Infrared Imaging System using secondary antibodies DyLight 680 or 800 (Invitrogen) at a dilution of 1:5000.

### Subcellular fractionation

Cells were resuspended in buffer A (10 mM HEPES (pH 7.9), 10 mM KCl, 1.5 mM MgCl2, 0.34 M sucrose, 10% glycerol, 1 mM DTT and 0.1%Triton X-100 containing 1 mM sodium vanadate and 1X Protein Inhibitor cocktail (Merck Life Sciences, P8340) and left for 5 min on ice. A fraction was collected and resuspended directly in sample buffer (TCL, total cell lysate). The rest of the cells were centrifuged at 4500 rpm for 5 min at 4 °C and the collected supernatant was considered the cytoplasmic fraction. Pellets were washed twice with buffer A and centrifuged at 4500 rpm for 5 min at 4°C. The final pellet was considered the nuclear fraction. All fractions were sonicated before analysis by SDS-PAGE

### Statistical analysis

All statistical analyses were performed using GraphPad Prism 9-11 software. Data are presented as mean ±standard error of the mean (SEM) unless otherwise indicated. To enable comparisons across independent experiments, the MFI data was normalized using the following procedure. The mean intensity of each individual experiment (*Ie*) and the overall mean intensity across all combined experiments (*Im*) were calculated. For each experiment, a normalization factor (*Im/Ie*) was computed and multiplied by the expression levels of all samples in that experiment.

Normally distributed data were analyzed using either a two-tailed unpaired t-test, or a one- or two-way ANOVA with Dunnet, Bonferroni or Tukey post hoc test for multiple comparisons. Non-normally distributed data were analyzed using a Kruskal Wallis-Dunn’s test or Mann-Whitney U test. P-value (p) was calculated and is expressed as *, p < 0.05; **, p <0.01; ***, p < 0.001 and ****, p <0.0001. p<0.05 was considered to be statistically significant.

## Acknowledgements

We thank Lidia Mateo and Camille Stephan-Otto Attolini (Biostatistics/Bioinformatics core facility) and the staff of the Functional Genomics, Mouse Mutant and Histopathology core facilities of IRB Barcelona, and the fluorescence-activated cell sorting Facility of UB-PCB for excellent technical support. We are very grateful to Maria Casanova (CNIO, Madrid) for useful suggestions and discussions on transplantation experiments, and to Luciano Di Croce and Cecilia Ballare (CRG, Barcelona) for technical insights with the ChIP experiments. This work was supported by grants to A.R.N. from the Fundación Científica de la Asociación Española Contra el Cancer (AECC, PROYE18035RODR), the Spanish Ministerio de Ciencia, Innovación y Universidades (MICIU, PID2019-109521RB-I00 funded by MCIN/AEI/10.13039/501100011033 and PID2022-136646OB-I00 funded by MICIU/AEI /10.13039/501100011033/FEDER, UE) and the Catalan Agency for Management of University and Research Grants (AGAUR, 2021 SGR-909), and to A.F.V. (MICIU PID2020-119875RB-I00). C.B.-E. was supported by the COFUND action of IRB Barcelona and BIST, which was funded by the European Union’s Horizon 2020 research and innovation programme under the Marie Skłodowska-Curie grant agreement No. 754558. We gratefully acknowledge institutional funding from IRB Barcelona, the CERCA Programme of the Catalan Government, and the MICINN through the Centres of Excellence Severo Ochoa award.

## Author contributions

M.C.-R. and C.B.-E. performed and designed experiments, analyzed the data, prepared the figures, and wrote the manuscript. E.I.R. performed mouse experiments. L.A.-B. performed cellular and molecular experiments. R.V. conducted and interpreted the histopathological analysis of HDM studies. E.L. and V.H.-M. provided help with mouse experiments. M.P. performed antigen presentation experiments. S.M.A.F. was involved in the design and generation of p38aBAC mice. A.R. provided essential reagents. S.K.W. provided Itgax-Cre mice, help with experimental design and revised the manuscript. M.T. supervised and interpreted the antigen presentation experiments. A.F.V. provided essential reagents, participated in the experimental design and interpretation of HDM studies and revised the manuscript. A.R.N. conceived the project, designed experiments, obtained funding and wrote the manuscript.

## Data availability

Source Data are provided with this paper. RNA-seq data have been deposited to the GEO database with accession number GSE273360 (https://www.ncbi.nlm.nih.gov/geo/query/acc.cgi?acc=GSE273360) and Single cell RNA-seq data in ArrayExpress database with accession number E-MTAB-14317 (https://www.ebi.ac.uk/biostudies/arrayexpress/studies/E-MTAB-14317?key=06752d08-08e3-4f99-8c91-63ed1e650830).

## Supplementary information

**Supplementary Fig. 1.**
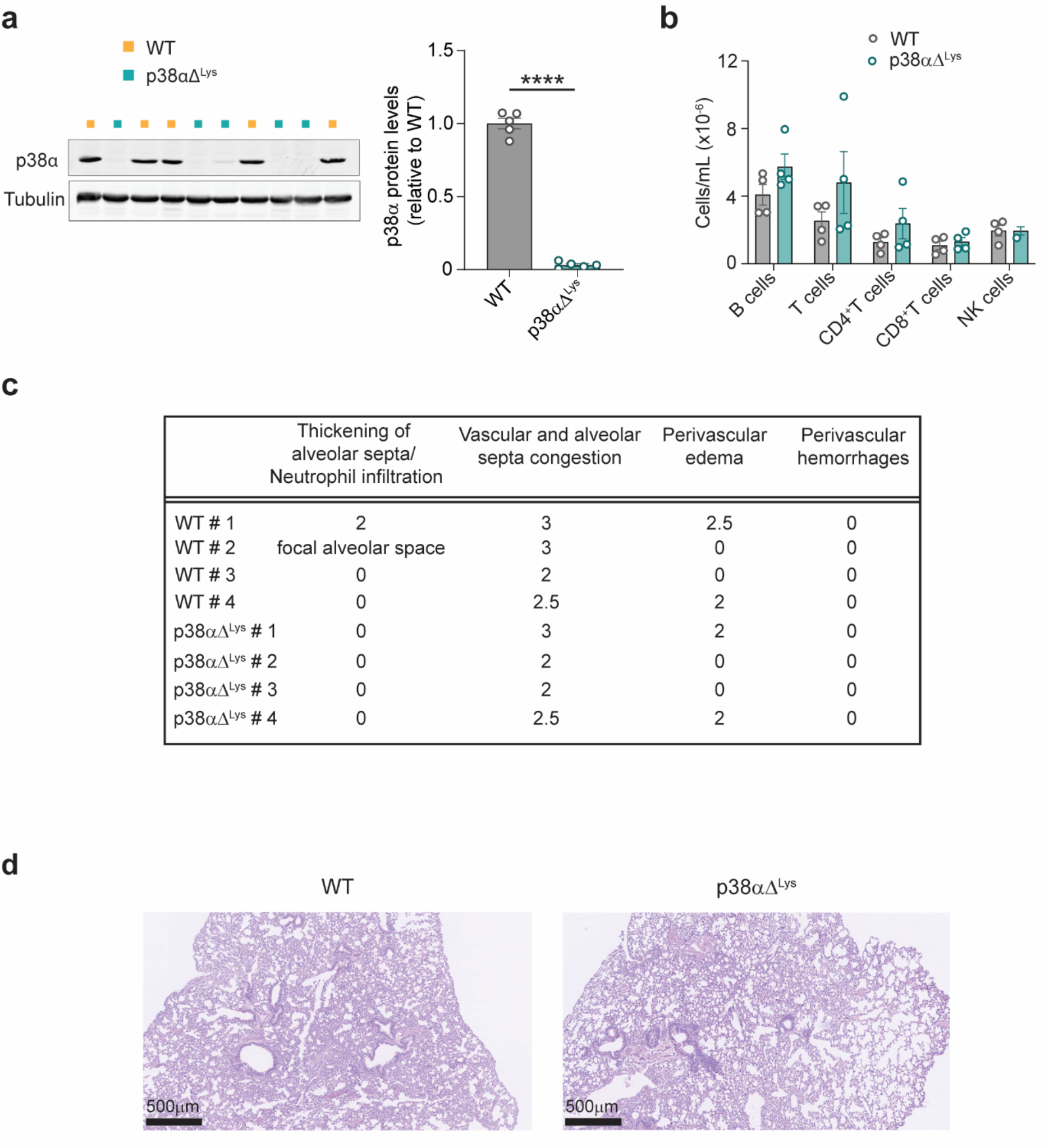
Characterization of p38a deletion in myeloid cells and the immune status of the lung. **a.** Immunoblotting of peritoneal macrophages extracted from WT and p38aΔ^Lys^ mice using the indicated antibodies. Histogram represents p38a/Tubulin ratios relative to WT mice. Each dot represents a cellular lysate obtained from peritoneal macrophages of a different mouse. Unpaired Student’s t test. ****, p<0.0001. **b.** Lymphoid cells from lungs of WT and p38aDLys mice were analyzed by FACS. Results are indicated as the total number of cells/mL in a suspension of 350 μL coming from one lobe of the lung. **c.** Histopathological analysis performed in a blinded fashion from hematoxylin and eosin staining of lungs from WT and p38aDLys mice. A semi-quantitative score was used to define the intensity of the inflammation: 0, none; 1, very mild; 2, mild; 3, moderate; 4, intense. Vascular and alveolar congestion might be due to the method of mice sacrifice more than to the inflammation *per se*. **d.** Representative H&E images from the lungs of WT and p38aDLys mice.

**Supplementary Fig. 2.**
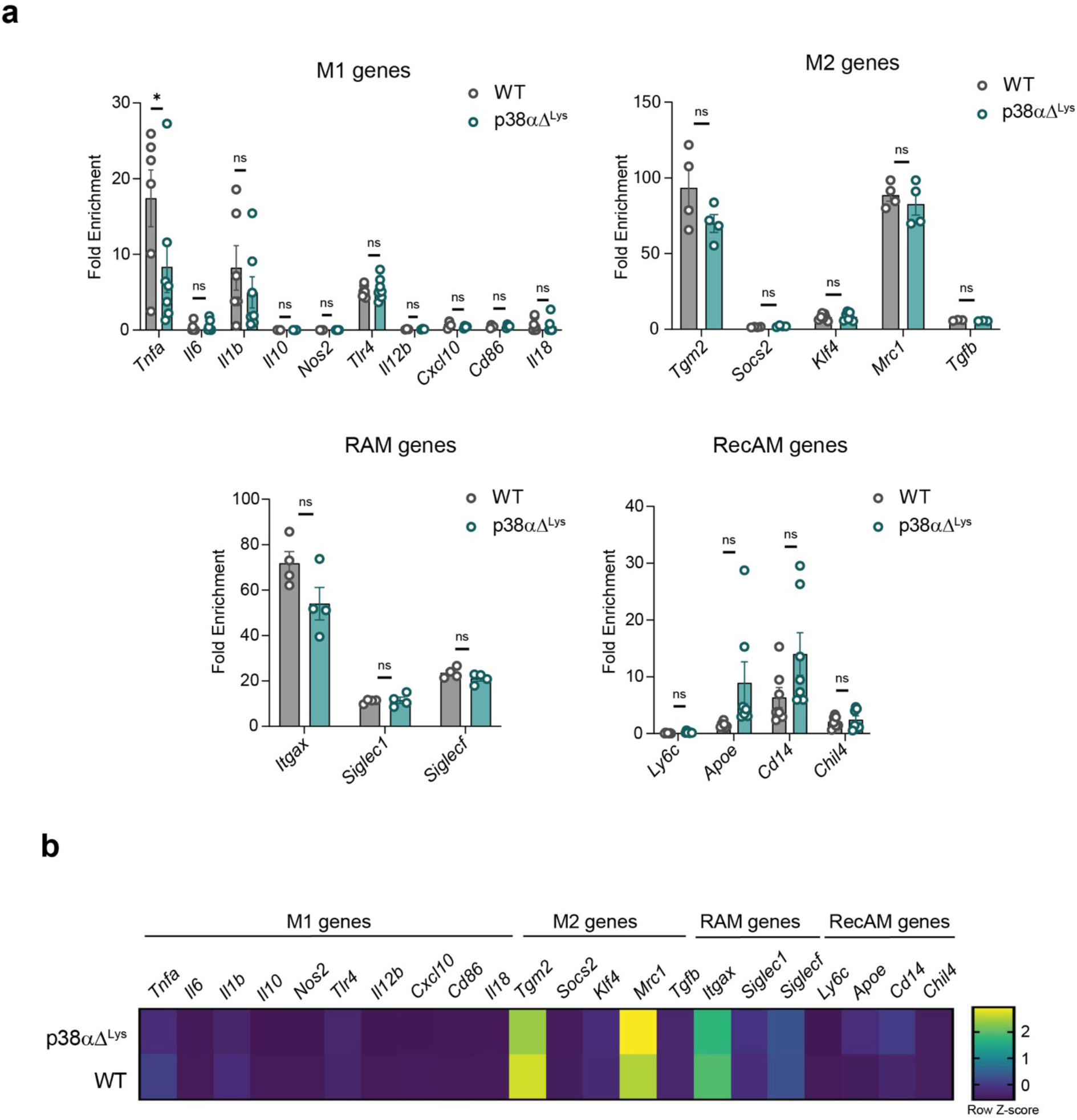
Expression analysis of M1, M2, RAM and RecAM genes in WT and p38aΔLys AMs. **a.** RT-qPCR analysis of mRNAs for the indicated genes in AMs obtained by BAL fluid from WT and p38aDLys mice. Data points represent the log-transformed difference between the target gene and the HPRT housekeeping gene to denote relative expression levels. Unpaired Student’s t-test to compare gene expression between genotypes. Data is shown as mean±SEM. *, p<0.05. ns, not significant. **b.** Heatmap showing the relative mRNA expression levels of the genes in (**a)** based on Z-score transformation.

**Supplementary Fig. 3.**
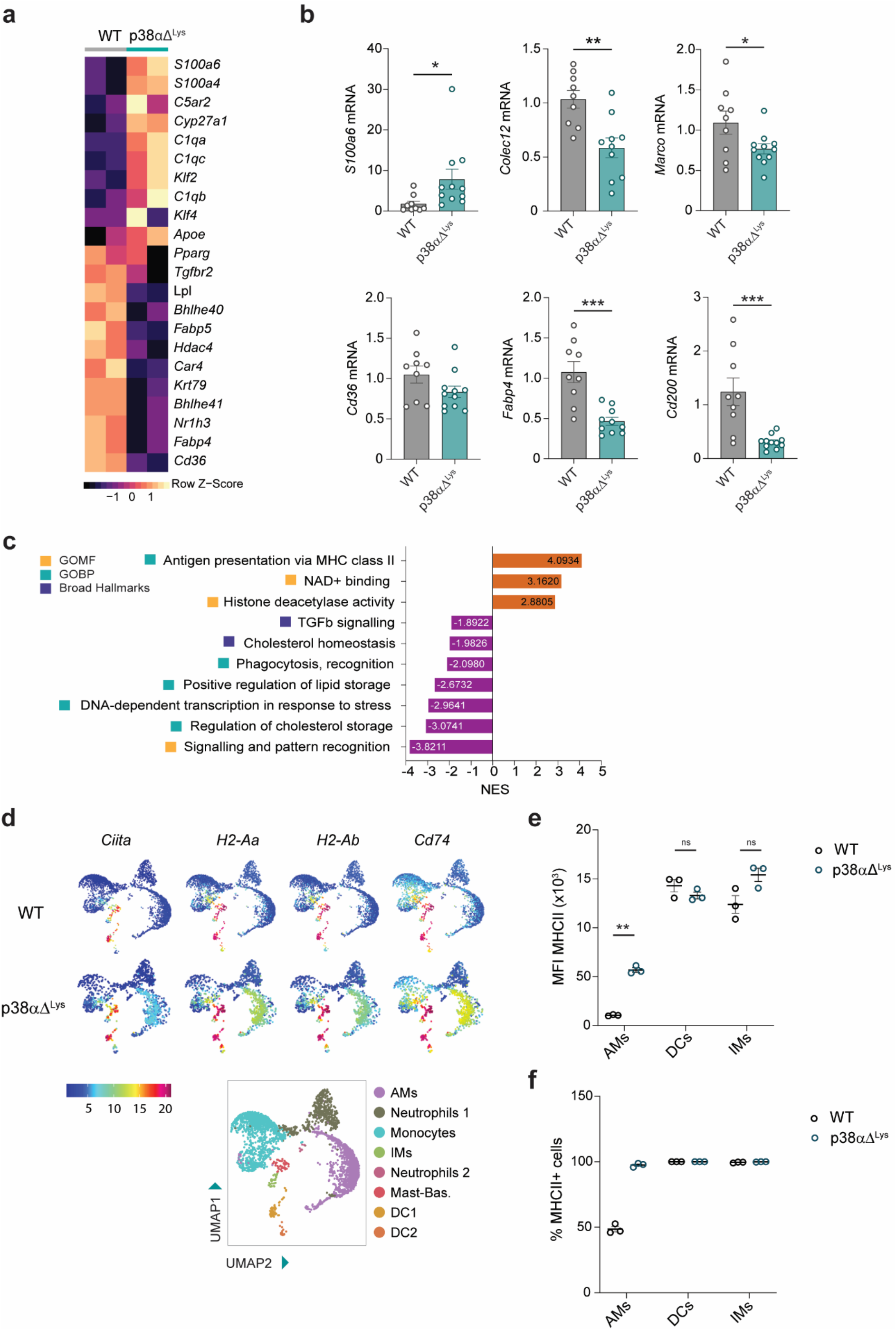
Upregulation of the antigen presentation machinery in p38α-deficient AMs. **a.** Heatmap representation of genes involved in different AM functions that were found significantly dysregulated in AMs from p38aDLys versus WT mice in the RNA-seq analysis. The expression levels are indicated by the color scale. p<0.05 for all the indicated genes. **b.** RT-qPCR analysis in AMs obtained from BAL fluid of WT and p38aDLys mice of representative genes found upregulated or downregulated in the RNA-seq analysis. Unpaired Student’s t test. Data shown as mean ±SEM. Each dot represents a different mouse. * p<0.05; ** p<0.01; *** p<0.001. **c.** Gene Set Enrichment Analysis (GSEA) analysis of the RNA-seq data showing selected GO terms enriched in AMs from p38aDLys mice. Normalized Enrichment Score (NES) values represented in graph bars are from the comparison of AMs from p38aDLys vs WT. **d.** Feature plots showing expression of the indicated genes in the myeloid sub-clustering populations of the scRNA-seq comparing WT and p38aDLys mice. The expression levels are indicated by the colors, and myeloid sub-clusters are identified in the legend. AMs=alveolar macrophages, IMs=interstitial macrophages, Mast-Bas=mastocyte-basophil cells, DC=dendritic cells. **e, f.** FACS analysis of MHCII expression in IMs (**e**) and DCs (**f**) from the whole lung of WT and p38aDLys mice. In **f**, histogram represents the percentage of MHCII^+^ cells for the indicated populations. Cells of each population were analyzed from the same individual mouse. Unpaired Student’s t test was used to compare the genotype effect in each population. **p<0.01.

**Supplementary Fig. 4.**
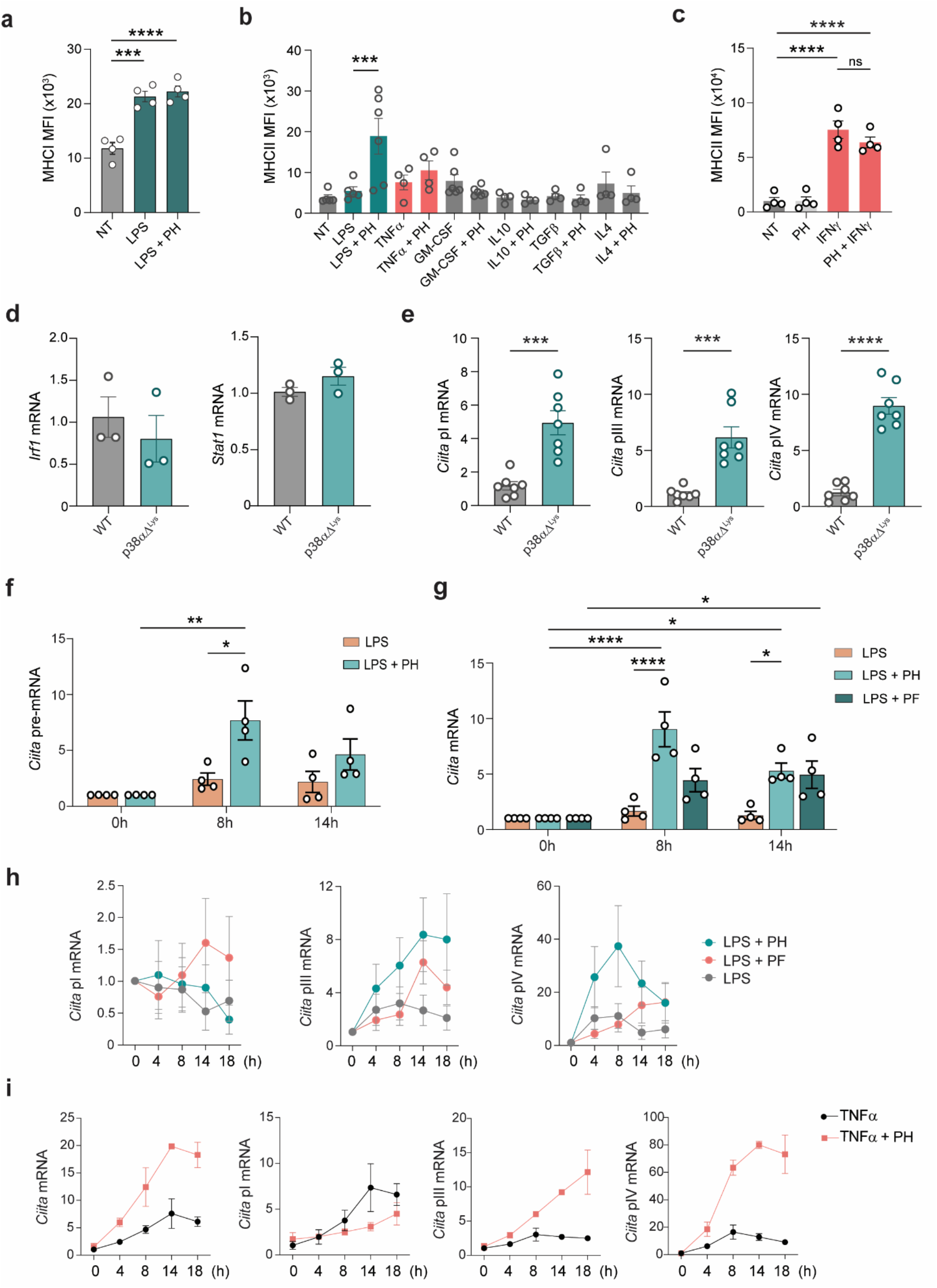
Mechanistic characterization of MHCII regulation by p38a. **a.** FACS analysis of MHCI expression in BMDMs pre-treated for 2 h with or without the p38a inhibitor PH797804 (PH) and then stimulated with LPS for 24 h. NT=non-treated control. One-way ANOVA test with a post hoc Dunnett’s test. Data is shown as mean±SEM. ***, p<0.001; ****, p<0.0001. **b.** FACS analysis of MHCII expression in BMDMs pre-treated for 2 h with or without the p38a inhibitor PH and then stimulated with LPS or the indicated cytokines for 24 h. NT=non-treated control. Each dot represents BMDMs from a different mouse. Data shown as mean ±SEM. One-way ANOVA test with a post hoc Dunnett’s test. ***, p<0.001. p-value refers to LPS condition. **c.** FACS analysis of MHCII expression in BMDMs pre-treated for 2 h with or without the p38a inhibitor PH and then stimulated with IFNγ for 24 h. Each dot represents BMDMs from a different mouse. One-way ANOVA test with a post hoc Bonferroni test. Data shown as mean±SEM. ****, p<0.0001. **d.** RT-qPCR analysis of IFNγ pathway targets in AMs obtained from BAL fluids of WT and p38aDLys mice. Each dot represents a different mouse. Data shown as mean ±SEM. **e.** RT-qPCR analysis of different *Ciita* transcripts in AMs obtained from BAL fluids of WT (n=7) and p38aDLys mice (n=7). Unpaired Student’s t test. Data shown as mean ±SEM. *** p<0.001; **** p<0.0001. **f.** RT-qPCR analysis of *Ciita* pre-mRNA in BMDMs pre-treated with or without the p38a inhibitor PH for 2 h and then stimulated with LPS for the indicated times. RNA was analyzed from BMDMs obtained from 4 different mice. Two-way ANOVA test with a post hoc Tukey test. Data represented as mean ±SEM. *, p<0.05; **; p<0.01. **g.** RT-qPCR analysis of *Ciita* mRNA in BMDMs pre-treated with or without inhibitors for p38a (PH) and MK2 (PF3644022, PF) for 2 h and then stimulated with LPS for the indicated times. RNA was analyzed from BMDMs obtained from four different mice. Two-way ANOVA test with a post hoc Dunnett’s test. Data represented as mean ±SEM. *, p<0.05; **; p<0.01; **** p<0.0001. **h.** RT-qPCR analysis of different *Ciita* transcripts in BMDMs pre-treated with or without inhibitors for p38a (PH) and MK2 (PF) for 2 h and then stimulated with LPS for the indicated times. RNA was analyzed from BMDMs obtained from three different mice. Data represented as mean ±SEM. **i.** RT-qPCR analysis of *Ciita* transcripts in BMDMs pre-treated with or without the p38a inhibitor PH for 2 h and then stimulated with TNFα for the indicated times. Each dot represents the median of two different mice. Data represented as median.

**Supplementary Fig. 5.**
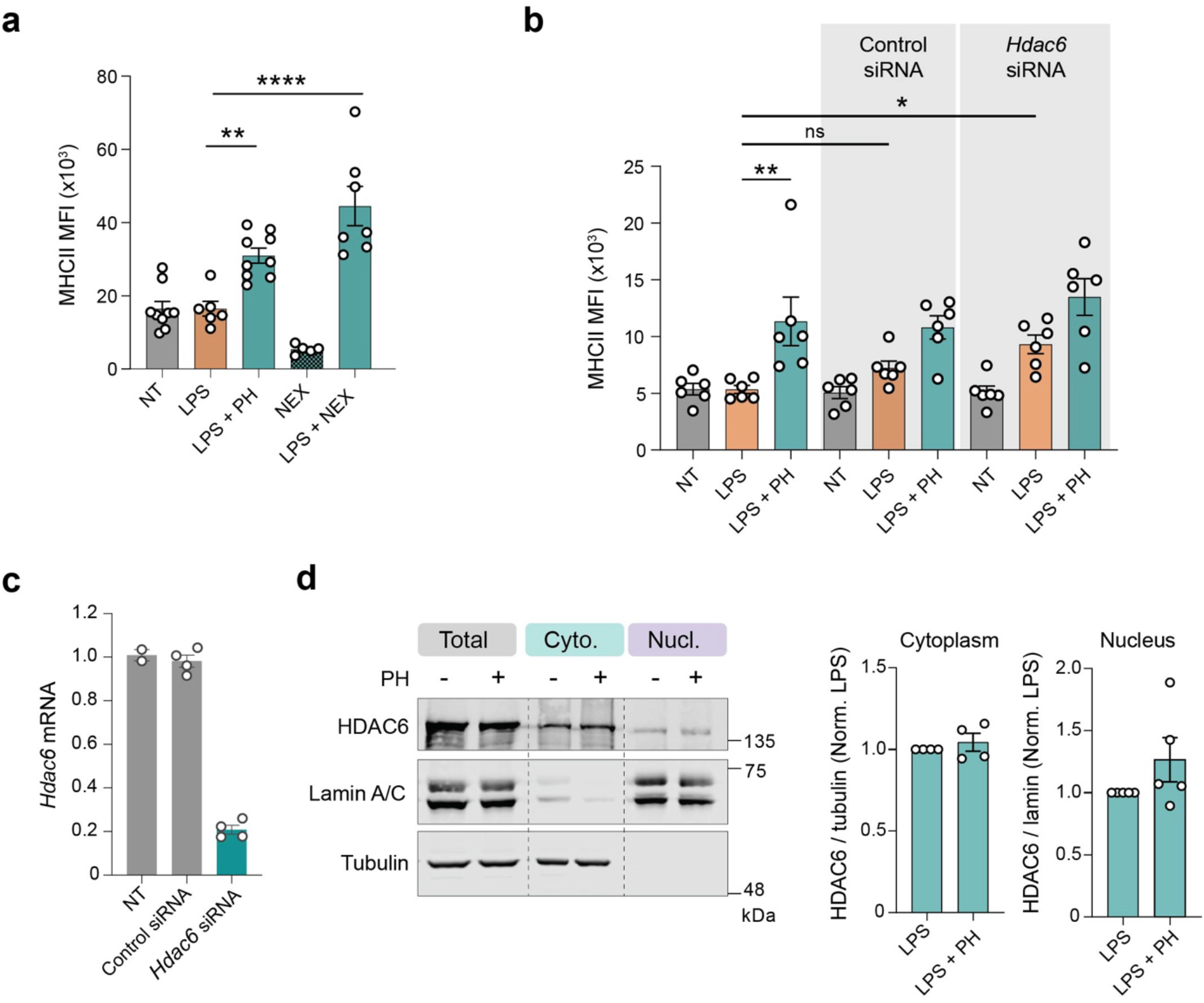
HDAC6 controls MHCII expression in macrophages. **a.** FACS analysis of MHCII expression in BMDMs treated for 24 h with LPS in the presence or absence of the p38a inhibitor PH797804 (PH) and the HDAC6 inhibitor Nexturastat A (NEX). NT=non-treated control. Each dot represents BMDMs from a different mouse. One-way ANOVA test with a post hoc Dunnett’s test. Data shown as mean ±SEM. p-val referred to LPS condition. **, p<0.01; ****, p<0.0001. **b.** FACS analysis of MHCII expression in BMDMs electroporated with control or HDAC6 siRNAs and then treated for 24 h with LPS in the presence or absence of the p38a inhibitor PH. NT=non-treated control. Data shown as mean ±SEM. Each point represents BMDMs from a different mouse. One-way ANOVA test with a post hoc Bonferroni test. Data shown as mean ±SEM. *, p<0.05; **, p<0.01. p-values refer to LPS condition. **c.** RT-qPCR analysis of *Hdac6* mRNA expression in BMDMs treated with control or *Hdac6* siRNAs. Data shown as mean ±SEM. Each point represents BMDMs from a different mouse. **d.** BMDMs were treated with or without the p38a inhibitor PH for 2 h and then stimulated with LPS for 2 h more. Cell lysates were separated into cytoplasm and nuclear fractions, which were analyzed by immunoblotting with the indicated antibodies. Tubulin was used as a cytoplasmic control and Lamin A/C as a nuclear control. Histograms represent the ratio HDAC6/Tubulin and HDAC6/Lamin in cytoplasmic and nuclear fraction, respectively, relative to LPS. Each point represents BMDMs from a different mouse.\

**Supplementary Fig. 6.**
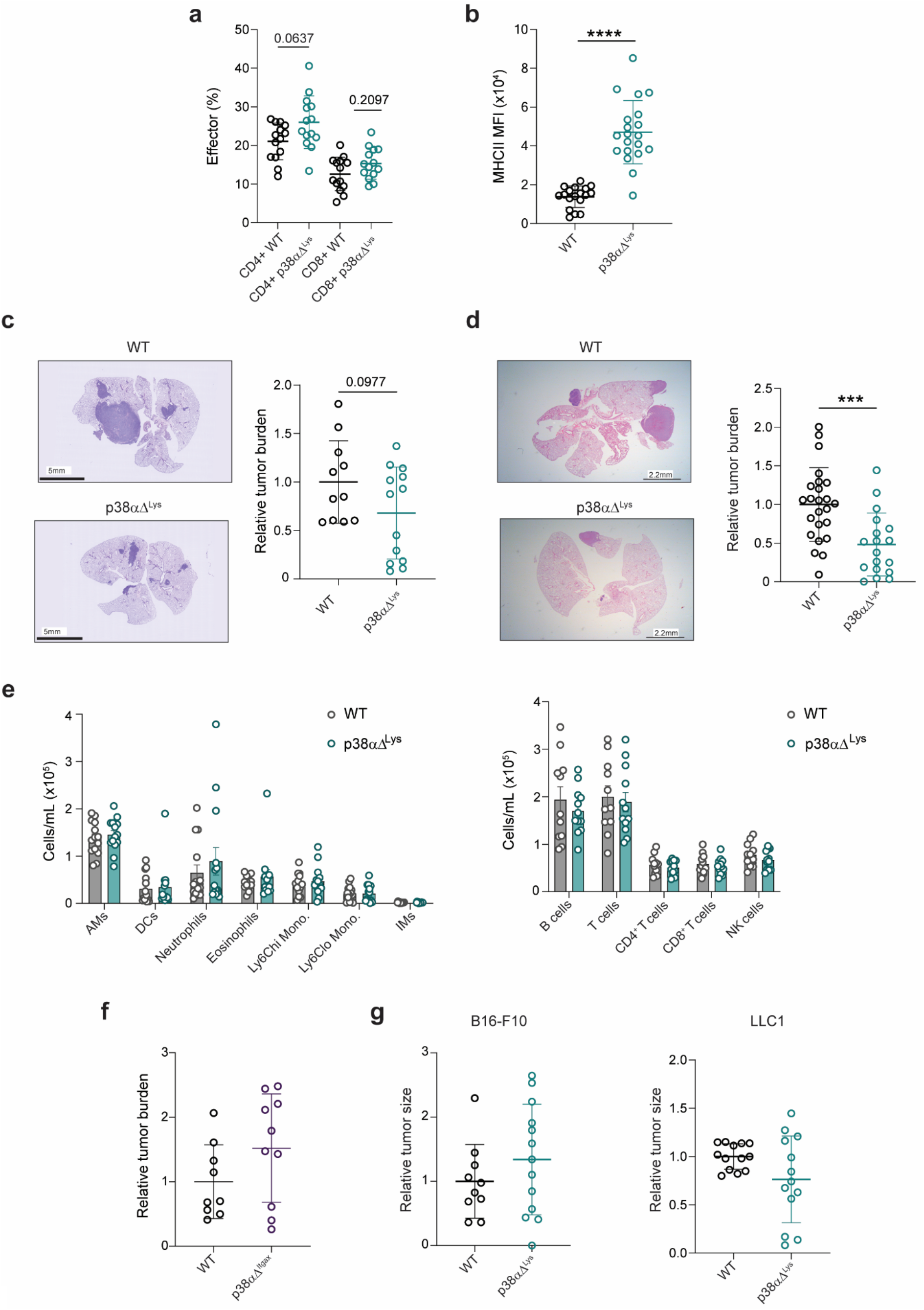
Impact of myeloid p38a on lung tumor growth and T cell activation. **a.** WT and p38aΔ^Lys^ mice were intravenously injected with B16-F10 cells and 21 days later the T cell activation profile in lungs was analyzed by FACS and represented as percentage of the total CD4^+^ and CD8^+^ T cells. Effector cells were defined as CD62L^low^, CD44^hi^. Mann Whitney test was used to compare the genotype effect in each population of T cells. Data shown as mean ±SEM (n=14) from two independent experiments. **b.** FACS analysis of MHCII expression in AMs from WT and p38aDLys mice after 21 days of B16-F10 cell inoculation. Data shown as mean from mice of 3 independent experiments. n=18-20. Unpaired Student’s t test. ****, p<0.0001. **c.** WT and p38aDLys mice were intravenously injected with B16-F10 cells and 21 days later lungs were harvested, fixed and H&E stained to quantify tumor burden (left panel). Quantification of the relative tumor burden for each genotype is shown on the right (right panel). Data are from three independent experiments (n=11-13 mice). Unpaired Student’s t test. Data shown as mean±SD. **d.** WT and p38aDLys mice were intravenously injected with LLC1 cells, and lung tumor burden was analyzed 17 days later. Lungs were harvested, fixed and H&E stained to quantify tumor burden (left panel). Data are from three independent experiments (n=18-24 mice). Data shown as mean±SD. Unpaired Student’s t test. ***, p<0.001. **e.** Myeloid cells (left) and lymphoid cells (right) from lungs with tumors of WT and p38aDLys mice were analyzed by FACS. Results are indicated as the total number of cells/mL in a suspension of 350 μl coming from one lobe of the lung. AMs=alveolar macrophages; DCs=dendritic cells; IMs=interstitial macrophages. **f.** WT and p38aDItgax mice were intravenously injected with B16-F10 cells and tumor burden was analyzed 21 days later. Data are from two independent experiments (n=9-10 mice). Data shown as mean±SD. **g.** WT and p38aDLys mice were subcutaneously injected with B16-F10 cells or LLC1 cells and tumor size was analyzed 21 days later. Data shown as mean±SD (n=10-13) from 2-3 independent experiments.

**Supplementary Fig. 7.**
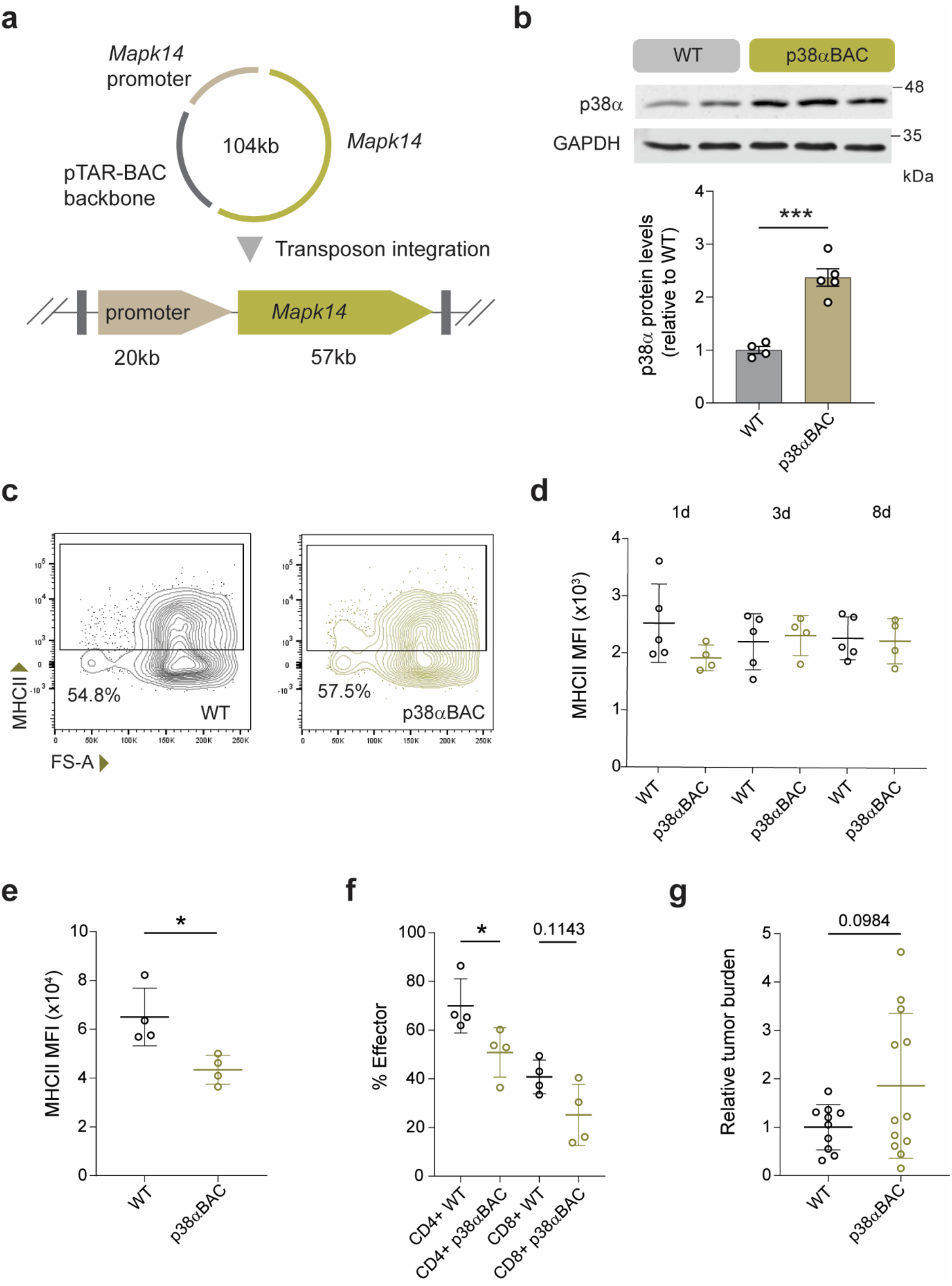
Characterization of p38aBAC mice during homeostasis and lung tumorigenesis. **a.** p38aBAC mice were generated to have additional copies of the p38α-encoding *Mapk14* gene integrated in the genome, using a Bacteria Artificial Chromosome (BAC) covering the entire *Mapk14* coding region plus 20 kb of upstream sequence. **b.** AMs were extracted from WT and p38aBAC mice and were analyzed by immunoblotting with the indicated antibodies. Histogram represents the p38a/GAPDH ratio relative to WT mice in cellular lysates obtained from BAL fluid. Unpaired Student’s t test. Data are presented as the mean ±SEM. ***, p<0.001. **c.** Representative FACS plots of MHCII analysis in AMs from WT and p38aBAC mice. **d.** FACS analysis of MHCII expression in AMs from WT and p38aBAC mice, which were cultured *ex vivo* for 1, 3 and 8 days. Each dot represents AMs from a different mouse (n=4). Data shown as mean ±SEM. **e.** WT and p38aBAC mice were inoculated with B16-F10 cells via tail vein and 21 days later MHCII expression in AMs was analyzed by FACS. Mean Fluorescence Intensity (MFI) is shown as mean ±SEM. Unpaired Student’s t test. *, p<0.05. **f.** WT and p38aBAC mice were inoculated with B16-F10 cells and 21 days later the T cell activation profile in lungs was analyzed by FACS and represented as percentage of the total CD4^+^ and CD8^+^ T cells. Effector cells were defined as CD62L^low^, CD44^hi^. Mann Whitney test was used to compare the genotype effect in each population of T cells. Data shown as mean ±SEM (n=4). *, p<0.05. **g.** Quantification of the tumoral area in the lungs from WT and p38aBAC mice analyzed 21 days after the injection of B16-F10 cells. Data shown as mean±SD (n=10-12).\

**Supplementary Fig. 8.**
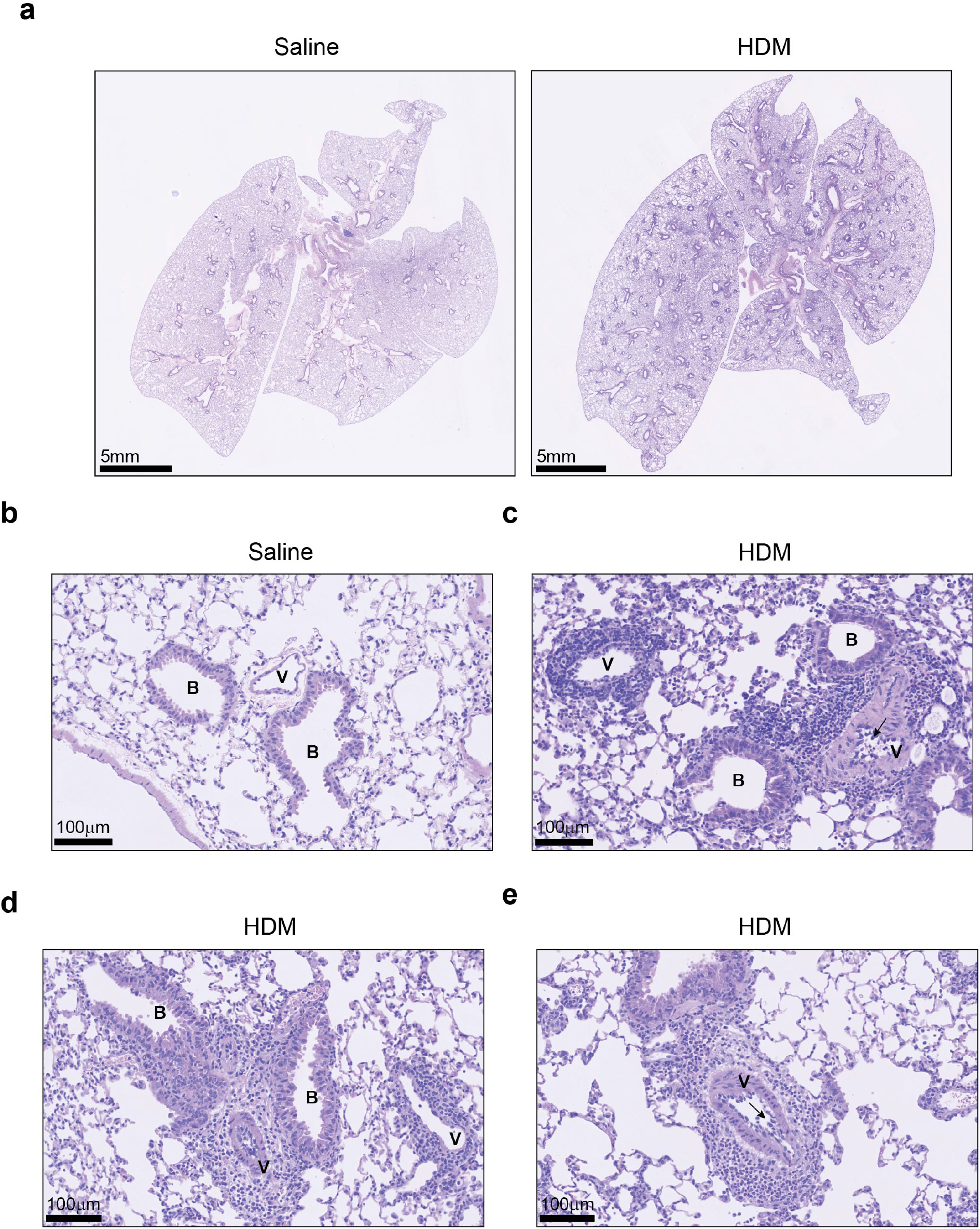
Histopathological analysis of the HDM model. **a.** Representative images of H&E-stained whole-lung sections from WT mice treated with control saline (left) or house dust mite (HDM) (right). Scale bars, 5 mm **b.** Higher magnification image of a H&E-stained lung section from a WT mouse treated with saline showing normal histology. Blood vessel (V); bronchiole (B). Scale bar, 100 μm. **c.** H&E-stained lung section from an HDM-exposed WT mouse showing the bronchovascular area with perivascular inflammation extending into the adjacent bronchioles, mild thickening of the tunica media and hypertrophic, activated endothelial cells (black arrow). Blood vessel (V); bronchiole (B). Scale bar, 100 μm. **d.** H&E-stained lung section from an HDM-exposed WT mouse showing the bronchovascular area with perivascular inflammation extending into the adjacent bronchioles and parenchyma. Mild epithelial cell hypertrophy in the bronchioles can also be observed. Blood vessel (V); bronchiole (B). Scale bar, 100 μm. **e.** H&E-stained lung section from a HDM-exposed WT mouse showing a blood vessel (V) with perivascular inflammation, a predominance of polymorphonuclear cells and fewer lymphocytes and macrophages. Activated, hypertrophic endothelial cells and focal infiltration of by inflammatory cells can also be observed (black arrow). Scale bar, 100 μm.

**Supplementary Fig. 9.**
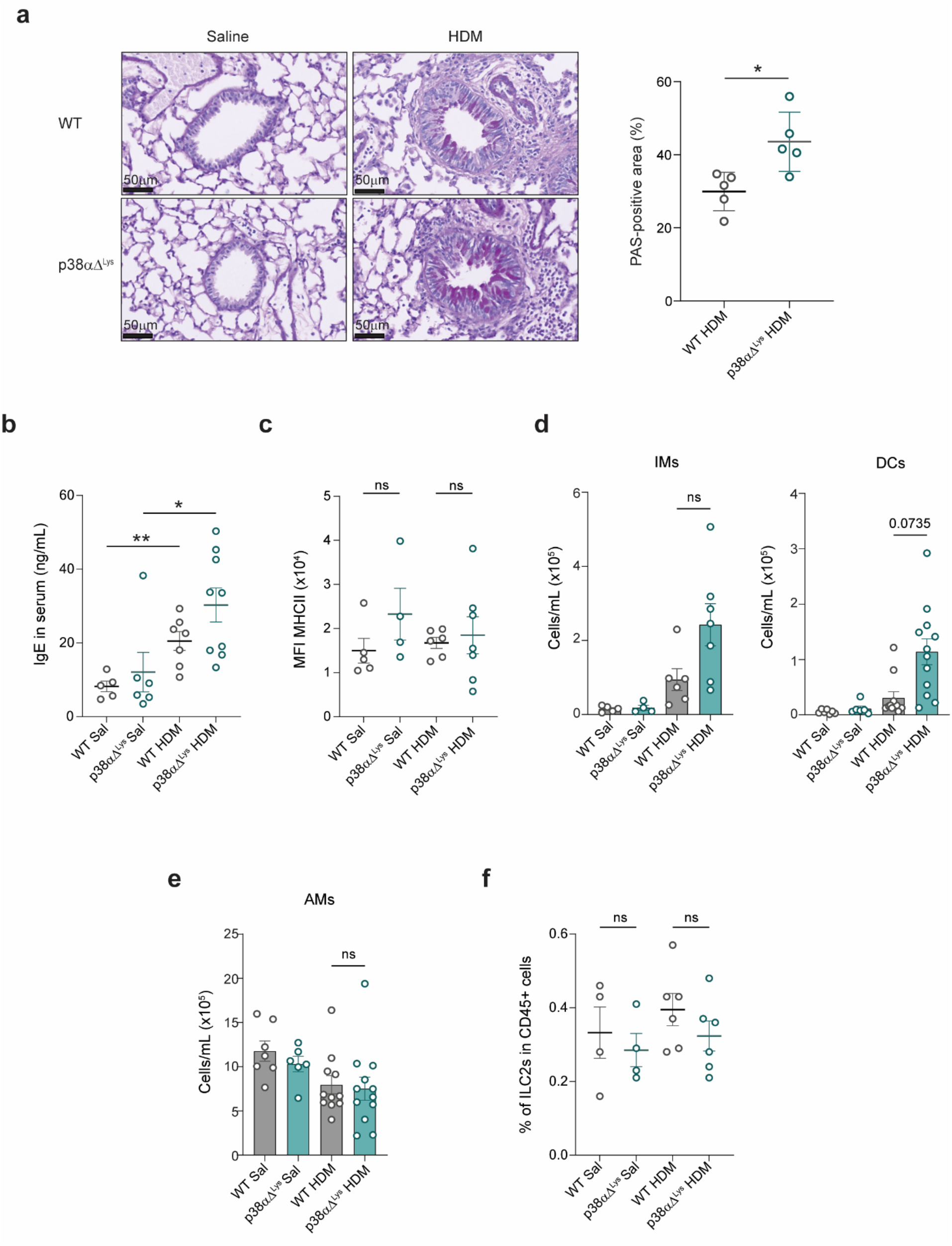
Characterization of the allergic airway inflammation in p38aΔ^Lys^ mice following HDM exposure. **a.** Representative Periodic acid-Schiff (PAS)-stained lung sections from WT and p38aDLys mice exposed to house dust mite (HDM) (left panel). Scale bar, 50 μm Quantification of the percentage of PAS-positive area is shown on the right (right panel). Mann Whitney test. *, p<0.05. **b.** Serum IgE levels were quantified by ELISA in mice exposed to HDM or saline solution. Unpaired Student’s t test was used to compare the treatment effect in each genotype. *, p<0.05; **<0.01. Data are presented as the mean ±SEM. n=5-9 from two independent experiments. **c.** FACS analysis of MHCII expression in interstitial macrophages (IMs) from BAL fluid of WT and p38aDLys mice after treatment with HDM allergen or saline solution (Sal) as a control. Each dot represents a different mouse. n= 4-7 from two independent experiments. **d-e.** Quantification of interstitial macrophages (IMs) and dendritic cells (DCs) (**d**) and alveolar macrophages (AMs) (**e**) in BAL fluid from WT and p38aDLys mice after treatment with HDM or saline solution. One-way ANOVA test with a post hoc Tukey’s test. Data is shown as mean±SEM. **f.** Quantification of ILC2 cell frequencies expressed as the percentage of CD45^+^ cells in mice treated with HDM or saline solution. ILC2 cells were defined as CD45^+^, Lin^-^ (B220, CD3, CD4, CD5, CD8α, CD11b, CD11c, CD19, Gr-1; TCRβ, Ter-119, γδTCR, NK1.1, FcεR1), CD90.2+ and ST2^+^. Data are presented as the mean ±SEM. n=4-6 mice from two independent experiments.

**Supplementary Fig. 10.**
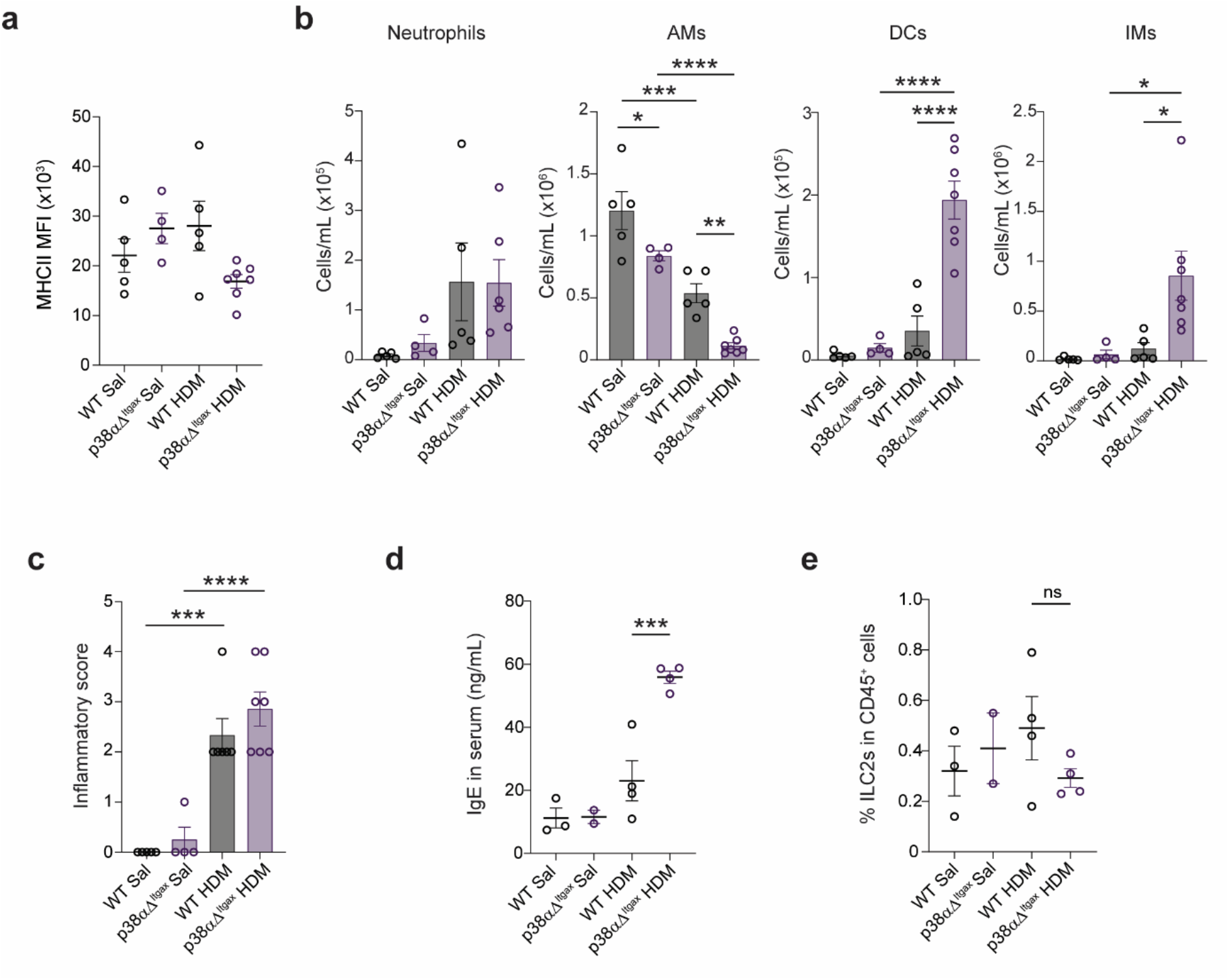
Allergic response in p38aΔ^Itgax^ mice upon HDM exposure. **a.** FACS analysis of MHCII expression in interstitial macrophages (IMs) from BAL fluid of WT and p38aDItgax mice after treatment with house dust mite (HDM) allergen or saline solution (Sal) as a control. **b.** Quantification of neutrophils, alveolar macrophages (AMs), dendritic cells (DCs) and IMs in BAL fluid from WT and p38aDItgax mice after treatment with HDM or saline solution. One-way ANOVA test with a post hoc Bonferroni’s test. Data is shown as mean±SEM. *, p<0.05; ***, p<0.001; ****, p<0.0001. **c.** Semiquantitative assessment of pulmonary inflammation. The indicated mice were treated with saline or HDM, and the severity grade was assigned in a blinded manner based on the severity and distribution of inflammation: grade 1 (minimal); grade 2 (slight); grade 3 (moderate); grade 4 (marked); grade 5 (severe). Data is shown as mean±SEM; ***, p<0.001. n=4-7 from two independent experiments. **d.** Serum IgE levels were quantified by ELISA in WT and p38aDItgax mice exposed to HDM or saline solution. One-way ANOVA test with a post hoc Bonferroni test. Data are presented as the mean ±SEM. ***, p<0.001. **e.** Quantification of ILC2 cell frequencies expressed as the percentage of CD45^+^ cells in mice treated with HDM or saline solution.

**Supplementary Fig. 11.**
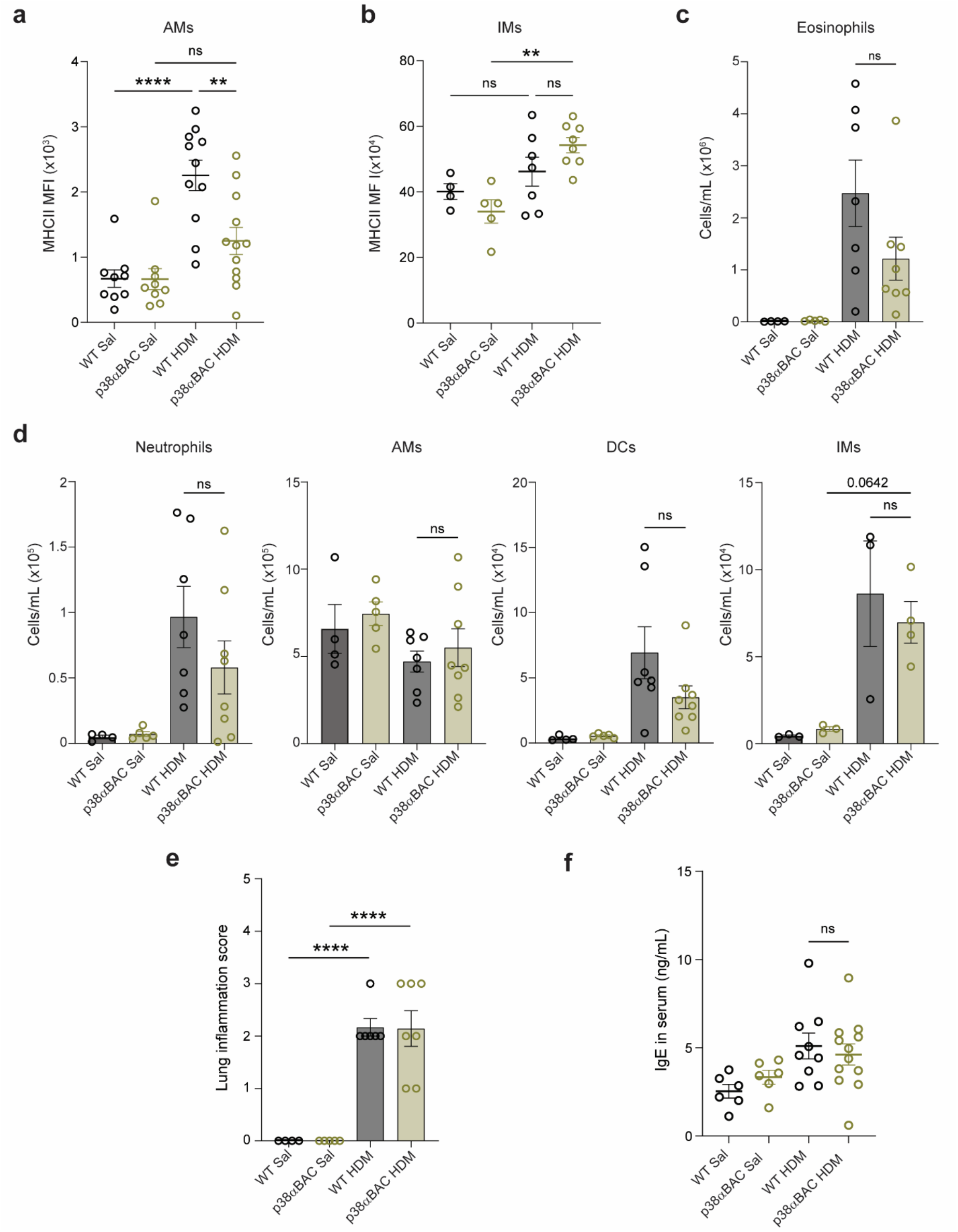
p38a overexpression attenuates allergen induced MHCII expression and allergic inflammation. a, **b.** FACS analysis of MHCII expression in alveolar macrophages (AMs) (**a**) or interstitial macrophages (IMs) (**b**) from WT and p38aBAC mice following exposure to house dust mite (HDM) allergen or saline solution (Sal) as a control. One-way ANOVA test with a post hoc Bonferroni test. Data shown as mean±SEM. **, p<0.01; ****, p<0.0001. n=8-12 from three independent experiments. **c, d.** Quantification of eosinophils (**c**) and the indicated myeloid cell populations (**d**) in BAL fluid from WT and p38aBAC mice after treatment with HDM or saline solution. One-way ANOVA test with a post hoc Bonferroni test. Data is shown as mean±SEM. n=4-8 from two independent experiments. **e.** Histological analysis of HDM-exposed lungs from WT and p38aBAC mice. The lung inflammatory score was calculated as described in Methods. One-way ANOVA test with a post hoc Bonferroni test. Data shown as mean±SEM. ****, p<0.0001. n=4-7 from two independent experiments. **f.** Serum IgE levels were quantified by ELISA in WT and p38aBAC mice exposed to HDM or saline solution. Data are presented as the mean ±SEM. n=6-12 from two independent experiments.

**Supplementary Fig. 12.**
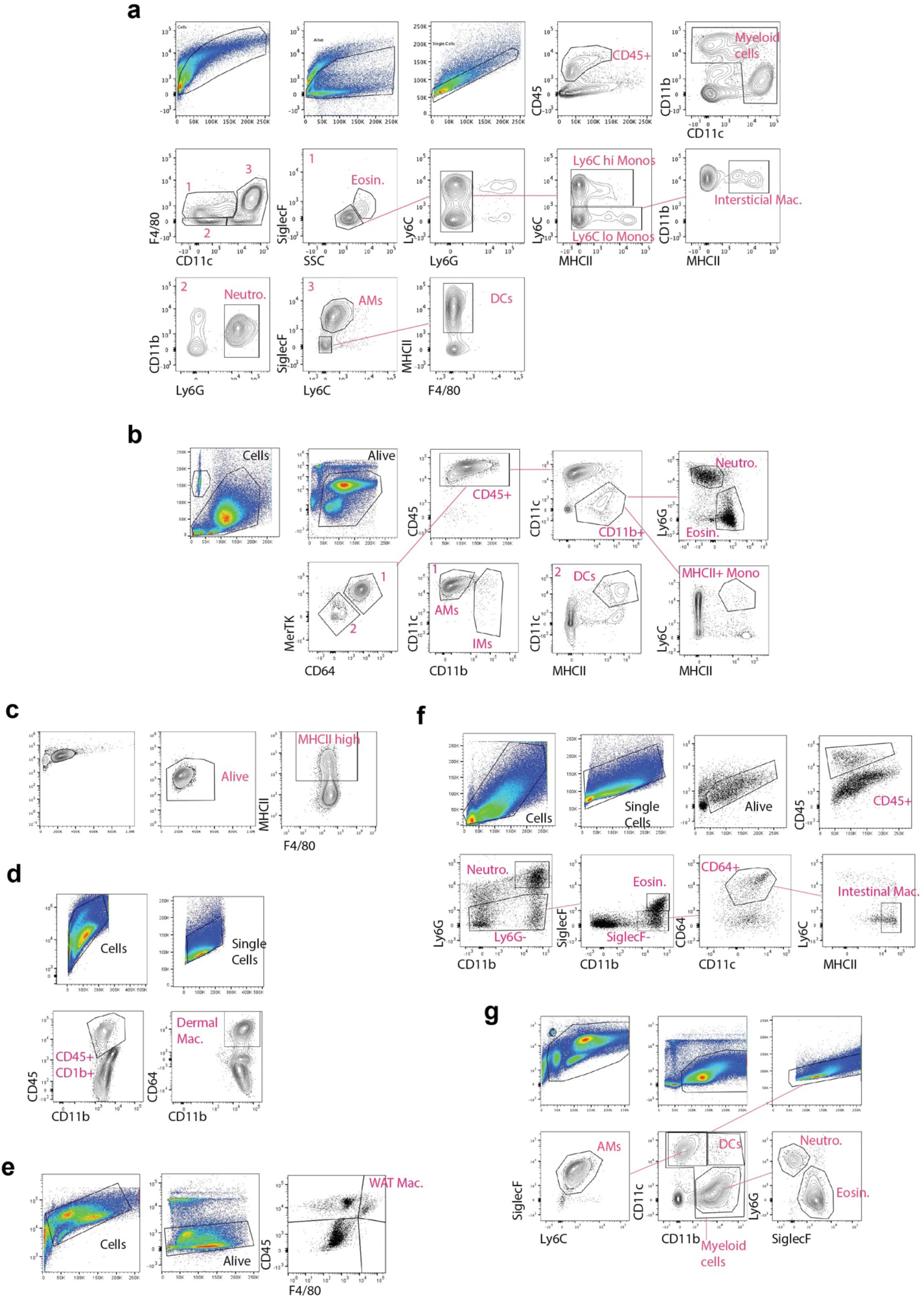
FACS gating for myeloid cell analysis. a, **b** FACS gating strategy for the myeloid cell populations in the lungs. Strategy (**b)** was used in BAL samples and included IMs in the analysis. **c.** FACS gating strategy for MHCII levels in macrophages. **d.** FACS gating strategy for sorting and analysis of dermal macrophages. **e.** FACS gating strategy for sorting and analysis of WAT macrophages. **f.** FACS gating strategy for sorting and analysis of intestinal macrophages. **g.** FACS gating strategy for myeloid populations in bronchoalveolar lavage (BAL).

**Supplementary Fig. 13.**
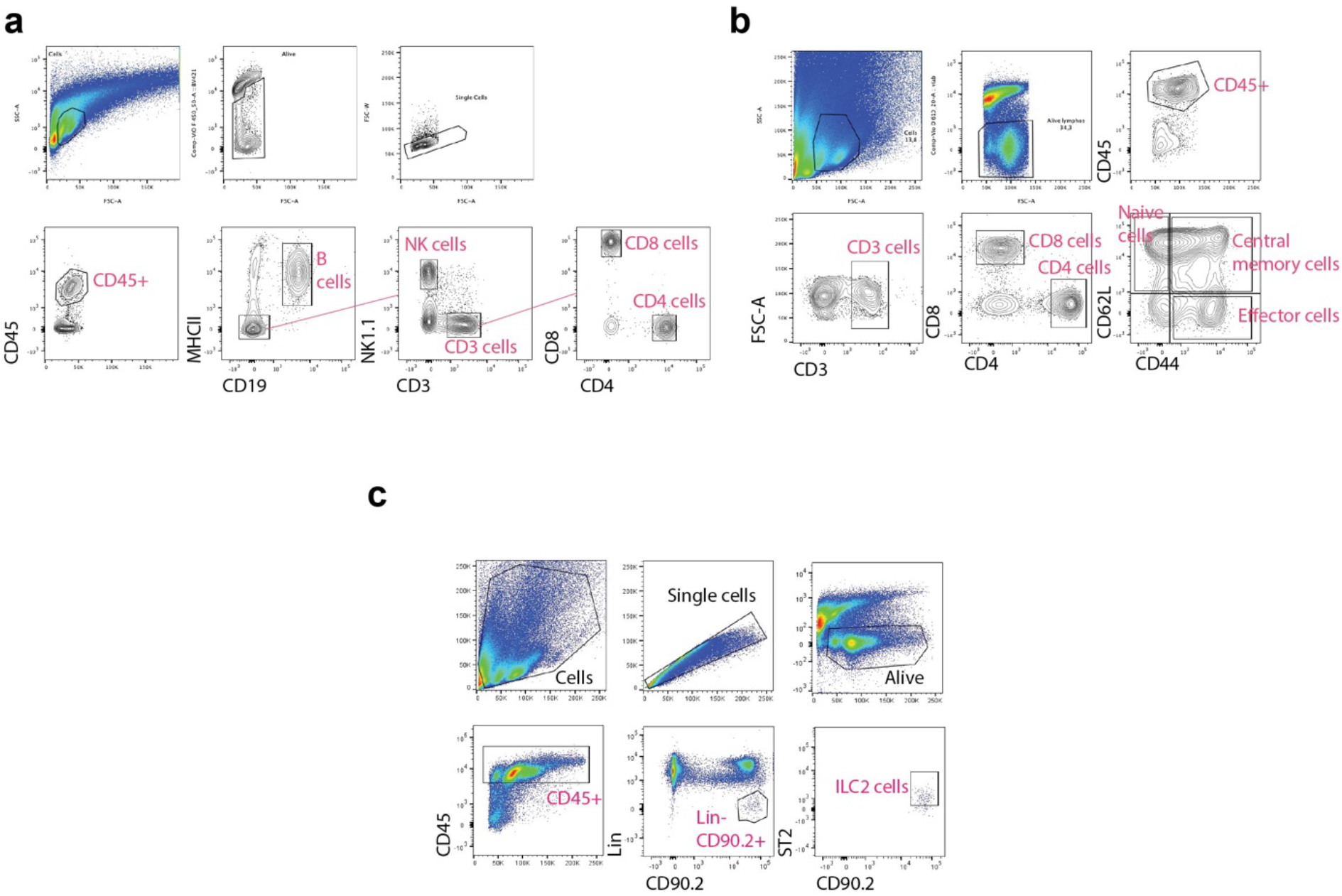
FACS gating for lymphoid cell analysis. **a.** FACS gating strategy for the lymphoid cell populations in the lungs. **b.** FACS gating strategy for the cell activation profiles of CD4 and CD8 T cells in the lungs. **c.** FACS gating strategy for the ILC2 cells in the lung.

**Supplementary Table 1.** Effect of epigenetic modulators on MHCII expression. BMDMs were treated with the indicated epigenetic drugs (Screen-Well Epigenetics Library BML-2836, Enzo) at the concentrations shown, and then stimulated with LPS for 24 h. MHCII expression was determined and the Values of Mean Fluorescence Intensity (MFI) were normalized to the treatment with LPS alone, which was given the value of 1 in all cases. ND = not done.

| **Number** | **Catalog number** | **Name** | **Activity** | **MHCII MFI at 2 µM** | **MHCII MFI at 10 µM** | **MHCII MFI at 20 µM** |
|---|---|---|---|---|---|---|
| 1 | GR309 | Trichostatin A | HDAC inhibitor | 1.04 | ND | ND |
| 2 | A283 | 2,4-Pyridinedicarboxylic Acid | Histone demethylase inhibitor |  | 0.74 | 0.97 |
| 3 | GR343 | Garcinol | HAT inhibitor |  | 1.09 | 3.87 |
| 4 | GR331 | Splitomicin | SIRT-2 inhibitor |  | 0.80 | 1.06 |
| 5 | GR330 | BML-210 | HDAC inhibitor |  | 1.63 | 4.58 |
| 6 | GR340 | Apicidin | HDAC inhibitor | 0.49 | ND | ND |
| 7 | GR323 | Suberoyl bis-hydroxamic acid | HDAC inhibitor | 1.24 | ND | ND |
| 8 | GR326 | Scriptaid | HDAC inhibitor | 1.15 | ND | ND |
| 9 | GR327 | Nullscript | Scriptaid Neg control |  | 0.46 | 0.71 |
| 10 | AC1526 | Decitabine | DNA Me transferase inhibitor |  | 0.90 | 1.03 |
| 11 | GR344 | Zebularine | DNA Me transferase inhibitor |  | 0.82 | 0.96 |
| 12 | 270288 | Vorinostat (SAHA) | HDAC inhibitor | 1.25 | ND | ND |
| 13 | AC1392 | Isonicotinamide | nicotinamide antagonist |  | 0.82 | 0.91 |
| 14 | GR350 | ITSA-1 | Inhibitor of TSA activity |  | 0.64 | 0.90 |
| 15 | EI320 | Sodium Phenylbutyrate | HDAC inhibitor |  | 0.94 | 0.81 |
| 16 | EI217 | Tranylcypromine hemisulfate | Lysine demethylase inhibitor |  | 1.09 | 0.92 |
| 17 | GR352 | Valproic acid | HDAC inhibitor |  | 0.78 | 0.99 |
| 18 | 270437 | 6-Chloro-2,3,4,9-tetrahydro-1H-carbazole-1-carboxamide | SIRT1 inhibitor |  | 0.92 | 1.04 |
| 19 | FR104 | Resveratrol | SIRT1 activator |  | 0.89 | 1.19 |
| 20 | AC1387 | M-344 | HDAC inhibitor | 1.45 | ND | ND |
| 21 | KI283 | Nicotinamide | SIRT inhibitor |  | 0.95 |  |
| 22 | GR346 | BML-266 | SIRT2 inhibitor |  | 1.51 |  |
| 23 | 270202 | Piceatannol | SIRT activator |  | 0.87 |  |
| 24 | GR353 | Fluoro-SAHA | HDAC inhibitor | 1.38 | ND | ND |
| 25 | GR351 | Valproic acid hydroxamate | HDAC inhibitor |  | 1.06 | 0.86 |
| 26 | 270484 | AGK2 | SIRT2 inhibitor |  | 0.83 | 3.72 |
| 27 | AC1388 | Salermide | SIRT inhibitor |  | 0.92 | 1.26 |
| 28 | 270344 | MC-1293 | HDAC inhibitor |  | 1.04 | 1.97 |
| 29 | 270381 | Anacardic acid | HAT inhibitor |  | 0.99 | 0.73 |
| 30 | 270485 | B2 | SIRT2 inhibitor |  | 0.74 |  |
| 31 | 270473 | BIX-01294 . Trichloride. Hydrate | Histone methyl transferase inhibitor |  | 0.97 | 1.98 |
| 32 | 270411 | Butyrolactone 3 | HAT inhibitor |  | 1.15 | 1.11 |
| 33 | 420033 | CTPB | HAT inhibitor |  | 0.73 | 0.97 |
| 34 | 270379 | Oxamflatin | HDAC inhibitor | 1.44 | ND | ND |
| 35 | 270308 | Sirtinol | SIRT inhibitor |  | 0.97 | 1.13 |
| 36 | G430 | Suramin. hexasodium salt | SIRT1 inhibitor |  | 0.98 | 0.77 |
| 37 | GR359 | BML-278 | SIRT1 actvator |  | 1.46 | 5.84 |
| 38 | AC1389 | NCH-51 | HDAC inhibitor | 2.89 | ND | ND |
| 39 | AC1390 | CI-994 | HDAC inhibitor |  | 2.24 | 10.52 |
| 40 | AC1391 | NSC-3852 | HDAC inhibitor | 0.84 | ND | ND |
| 41 | AC1566 | Aminoresveratrol sulfate | SIRT1 activator |  | 0.85 | 1.15 |
| 42 | GR361 | BML-281 | HDAC-6 inhibitor | 0.99 | 0.44 | ND |
| 43 | FR119 | Triacetylresveratrol | SIRT1 activator |  | 0.87 | 1.08 |

**Supplementary Table 2.**
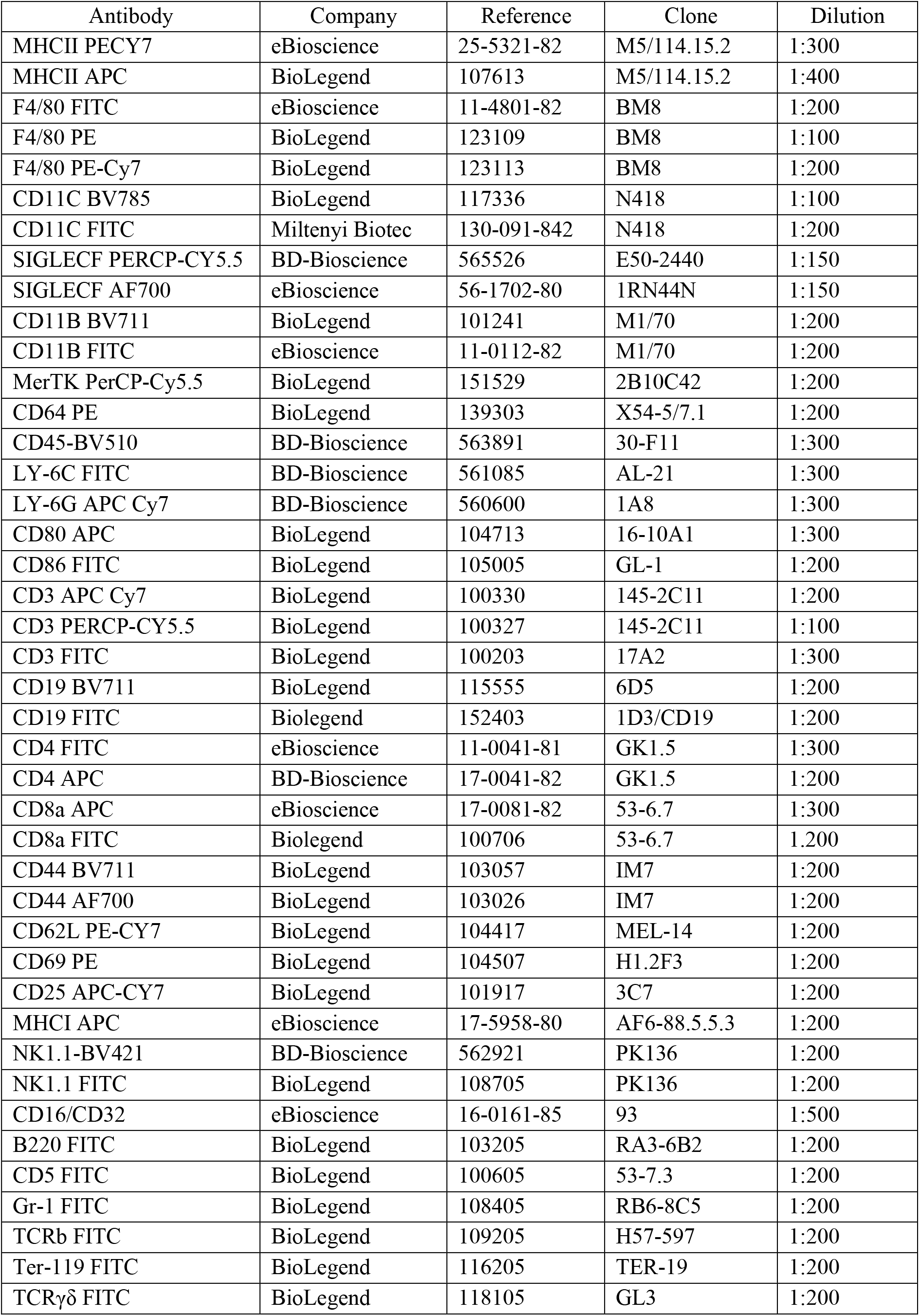
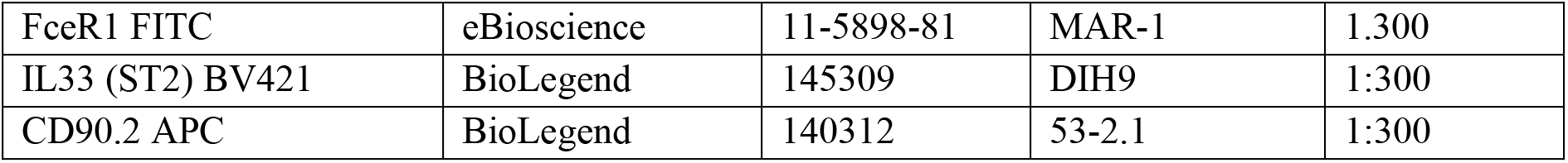
Antibodies used for FACS analysis.

**Supplementary Table 3.**
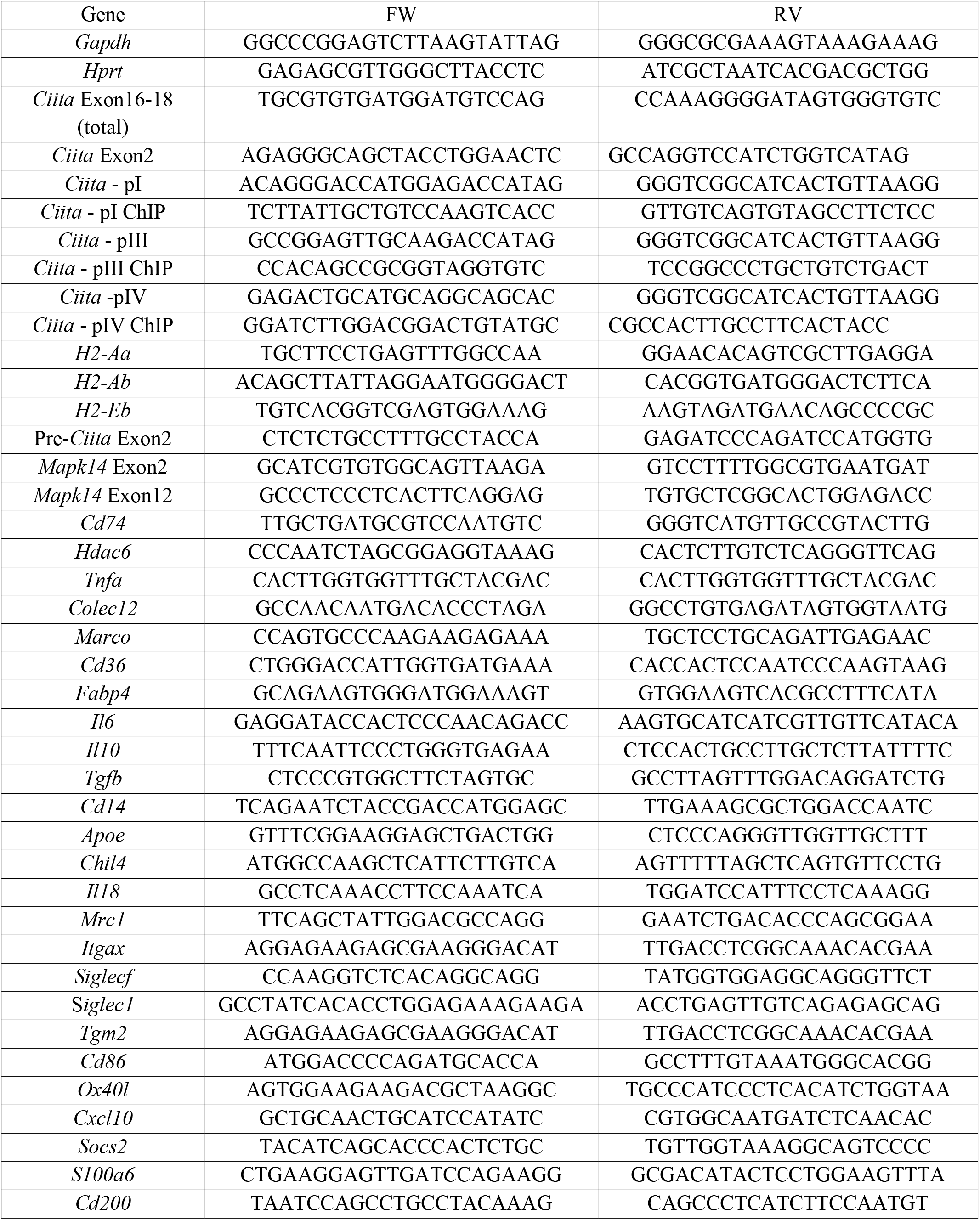
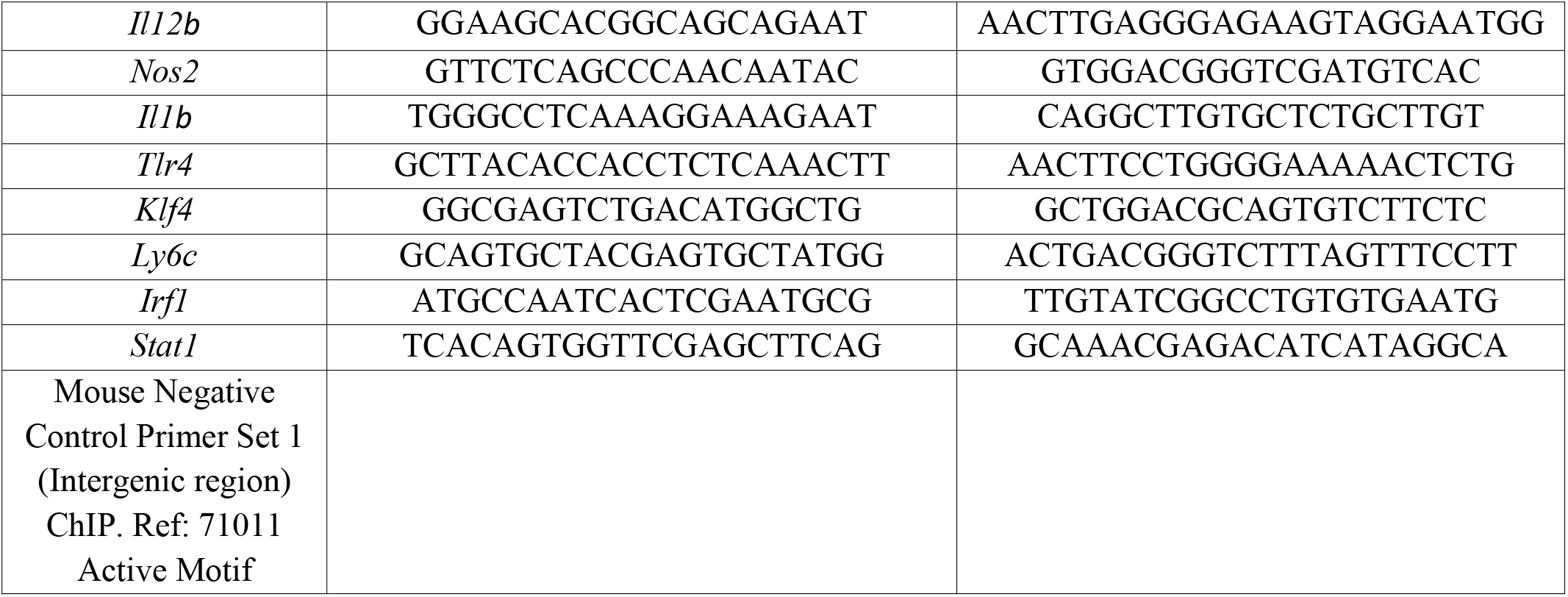
Primers used for RT-qPCR.

